# Physical and functional interaction between SET1/COMPASS complex component CFP-1 and a Sin3 HDAC complex

**DOI:** 10.1101/436147

**Authors:** F. Beurton, P. Stempor, M. Caron, A. Appert, Y. Dong, R. Chen, D. Cluet, Y. Couté, M. Herbette, N. Huang, H. Polveche, M. Spichty, C. Bedet, J. Ahringer, F. Palladino

## Abstract

The CFP1 CXXC zinc finger protein targets the SET1/COMPASS complex to non-methylated CpG rich promoters to implement tri-methylation of histone H3 Ly4 (H3K4me3). Although H3K4me3 is widely associated with gene expression, the effects of CFP1 loss depend on chromatin context, so it is important to understand the relationship between CFP1 and other chromatin factors. Using a proteomics approach, we identified an unexpected link between *C. elegans* CFP-1 and a Rpd3/Sin3 histone deacetylase complex. We find that mutants of CFP-1, SIN-3, and the catalytic subunit SET-2/SET1 have similar phenotypes and misregulate common genes. CFP-1 directly binds SIN-3 through a region including the conserved PAH1 domain and recruits SIN-3 and the HDA-1/HDAC subunit to H3K4me3 enriched promoters. Our results reveal a novel role for CFP-1 in mediating interaction between SET1/COMPASS and a Sin3 HDAC complex at promoters and uncover coordinate regulation of gene expression by chromatin complexes having distinct activities.

## Introduction

The CFP1/CXXC zinc finger protein targets the SET1/COMPASS complex to non-methylated CpG rich regions for trimethylation of histone H3 on Lys4 (H3K4me3) (Brown et al. 2017; Clouaire et al. 2012; Lee and Skalnik 2005; Mahadevan and Skalnik 2016; Thomson et al. 2010), a modification widely associated with active promoters (Bernstein et al. 2005; Heintzman et al. 2007; Schneider et al. 2004). The roles of CFP1 and the SET1/COMPASS complex in gene regulation are unclear. In different systems, loss of individual subunits does not have widespread effects on transcription, with only small subsets of genes affected (Clouaire et al., 2012, 2014; Howe et al., 2017; Lenstra et al., 2011; Margaritis et al., 2012). The effects vary depending on context, consistent with potential interactions with other factors and proposals that H3K4me3 may promote transcriptional memory and consistency (Howe et al. 2017).

In yeast, SET1 acts in a single complex known as COMPASS (complex of proteins associated with Set1) that is responsible for all H3K4 methylation (Miller et al. 2001; Roguev et al. 2001). In mammals by contrast, six complexes have been isolated defined by the catalytic subunits SET1A, SET1B, MLL1, MLL2, MLL3 and MLL4 (reviewed in (Shilatifard 2012). The enzymatic activity of SET1/MLL family members is regulated by interactions with additional proteins, including Swd3/WDR5, Swd1/RbBP5, Bre2/ASH2, and Sdc1/hDPY30 that influence the state (mono-, di-, or tri) of methylation deposited (Dehe et al., 2006; Dou et al., 2006; Steward et al., 2006). In addition, unique subunits including CFP1 are associated with each complex and contribute to its specificity (Hughes et al. 2004; Lee and Skalnik 2005; Lee et al. 2006; Narayanan et al. 2007; Tyagi et al. 2007).

SET1/MLL complexes have non-redundant functions, as demonstrated by the distinct phenotypes and embryonic lethality caused by deletion of individual SET1/MLL genes (Bledau et al. 2014; Glaser et al. 2006; Lee et al. 2008; Yu et al. 1995). While SET1 proteins are responsible for global H3K4me3 at promoter regions in different organisms (Ardehali et al. 2011; Hallson et al. 2012; Li and Kelly 2011; Wu et al. 2008; Xiao et al. 2011), MLL proteins deposit H3K4 methylation at specific genes or regulatory elements (Denissov et al. 2014; Hu et al. 2013; Wang et al. 2009a).

*Caenorhabditis elegans* contains a single homologue of SET1, named SET-2, one MLL-like protein, SET-16, and single homologs of WDR5, ASH2L, DPY30, RbBP5 and CFP1 (Li and Kelly 2011; Simonet et al. 2007; Xiao et al. 2011), simplifying functional studies of SET1/MLL regulatory networks. Inactivation of SET-2, WDR-5.1, DPY-30, RbBP5 and CFP-1 has shown that they all contribute to global H3K4 methylation in the germline and soma, and share common functions in somatic and germline development (Greer et al., 2010; Li and Kelly, 2011; Robert et al., 2014; Simonet et al., 2007; Xiao et al., 2011; Xu and Strome, 2001; Han et al., 2017). To biochemically analyze the complex and identify associated proteins that may contribute to its functional outcome, we immunoprecipitated tagged CFP-1 and WDR-5.1, and identified copurifying proteins by mass spectrometry. In addition to identifying distinct SET-2/SET1 and SET-16/MLL complexes, we found that WDR-5.1 co-immunoprecipitates NSL histone acetyltransferase (HAT) complex subunits (Cai et al., 2010; Dias et al., 2014; Raja et al., 2010; Zhao et al., 2013). Most importantly, we show that CFP-1 physically and functionally interacts with a conserved Rpd3/Sin3 histone deacetylase complex. Mutants of SET-2/SET1 and Rpd3/Sin3 complex subunits share common phenotypes, and CFP-1 is important for recruitment of both SIN-3 and the HDA-1/HDAC subunit to H3K4me3 enriched promoters. Our results reveal a novel role for CFP-1 in bridging interactions between the SET-2/SET1 and Rpd3/Sin3 HDAC complexes to maintain the embryonic transcriptional program and influence both somatic and germline development.

## Results

### Co-immunoprecipitation of subunits of the *C. elegans* SET1/COMPASS complex

We used a proteomics approach to characterize *C. elegans* COMPASS-like complexes and search for associated proteins. In addition to the catalytic subunit, SET1/COMPASS complexes contain the core components ASH2, RbBP5, WDR5 and DPY30, and the unique subunits CFP1/CXXC and WDR82 (Cosgrove and Patel, 2010; Lee and Skalnik, 2005, 2008; Lee et al., 2007; Steward et al., 2006; Takahashi et al., 2011; Wu et al., 2008). Using strains containing two previously described transgenes, CFP-1::GFP and HA::WDR-5.1 (Chen et al. 2014; Simonet et al. 2007), we found that CFP-1::GFP coprecipitated HA::WDR-5.1, and that HA::WDR-5.1 coprecipitated CFP-1::GFP (Figure 1A and S2A). In addition, probing individual precipitates with anti-ASH-2 and anti-DPY-30 antibodies revealed the presence of both proteins. Because single or low copy tagged SET-2 protein could not be detected, we were unable to confirm its presence in the IP experiments.

**Figure 1.**
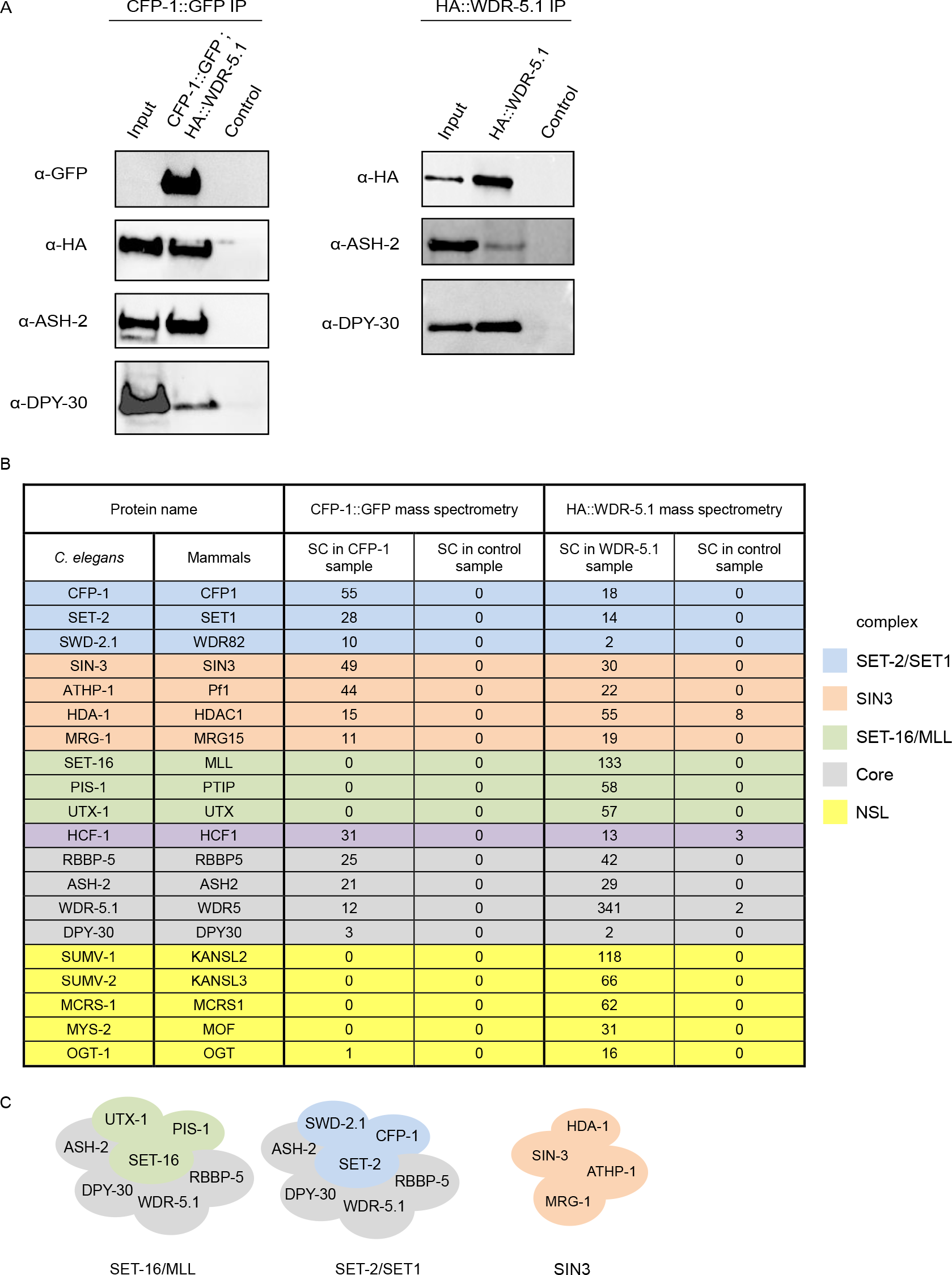
The conserved SIN3 complex copurifies with CFP-1 and WDR-5.1. (A) Co-IP of CFP-1, WDR-5.1, ASH-2 and DPY-30. Western blot analysis of CFP-1::GFP (left panel) and HA::WDR-5.1 (right panel) purified complexes from embryos. For CFP-1::GFP IP, immunoprecipitations were performed on 4 mg total protein extract, and loadings were 1/200 of total protein extract for input and 1/2 of total elution volume. For HA::WDR-5.1 co-IP, elutions used for mass spectrometry analysis were probed with anti-ASH-2 or anti-DPY-30 antibodies. Loadings were: 1/8000 of total protein extract for input and 1/250 of elution for ASH-2; 1/60 000 of total protein extract for input and 1/200 of elution for DPY-30. The anti-HA membrane corresponds to the one in Figure S2B. (B) List of selected proteins identified by mass spectrometry of CFP-1::GFP or HA::WDR-5.1 immunoprecipitations, and their mammalian homologue. Subunits specific to the SET-2/SET1, SET-16/MLL, SIN3 and NSL complexes are highlighted in blue, green, orange and yellow, respectively. SET1/MLL core complex subunits are highlighted in grey. HCF-1 copurifies with both SET-2/SET1 and SET-16/MLL complexes, but is not a core complex subunit. SC; Spectral Counts. In WDR-5.1 mass spectrometry, 242 proteins were found with a spectral count (SC) WDR-5.1 ≥ 3 and a SC control = 0 or SC WDR-5.1/SC control ≥ 5 for at least one replicate; in CFP-1 mass spectrometry 178 proteins were found with a SC CFP-1 ≥ 3 and a SC control = 0. (C) Cartoon representation of SET-2/SET1, SET-16/MLL and SIN3 complexes; subunits are highlighted as in (B).

The above experiments show that in embryos, CFP-1::GFP and HA::WDR-5.1 tagged proteins associated with each other *in vivo* and co-immunoprecipitate native ASH-2 and DPY-30, consistent with their incorporation into a SET1-related complex. To define SET1/MLL complexes and identify additional associated proteins, we undertook mass spectrometry-based proteomic characterization of CFP-1::GFP and HA::WDR-5.1 immunoprecipitates. We reasoned that WDR-5.1, a core component of SET1/MLL complexes, should immunoprecipitate both SET1 and MLL-related complexes (Dou et al., 2006; van Nuland et al., 2013; Patel et al., 2009), while CFP-1 should specifically immunoprecipitate SET1, but not MLL-related complexes (Clouaire et al. 2012; Lee and Skalnik 2005). Both tagged proteins were detected as unique bands in immunoprecipitates obtained using either anti-GFP or anti-HA antibodies (Fig. S2B), and as predominant bands by silver-staining (Fig. S2C). Tandem mass spectrometry (MS/MS) based proteomic analyses of the immunoprecipitates and comparison with eluates from negative controls identified both common and unique binding partners of CFP-1 and WDR-5 (Figures 1B; see Table S1 for a full list). We found that CFP-1::GFP and HA::WDR-5.1 immunoprecipitates contained all common subunits of SET1/MLL complexes, including Swd1/RBBP-5, Bre2/ASH-2, and Sdc1/DPY-30. The orthologue of human host cell factor HCF1, a transcriptional regulator associated with COMPASS-like and other chromatin-associated complexes (Zargar and Tyagi 2012) was also identified in both immunoprecipitates. WDR-5.1 additionally coprecipitated specific components of an MLL-related complex including the MLL-like histone methyltransferases SET-16, the histone H3K27 demethylase UTX-1, and PIS-1 (Vandamme et al. 2012). Conversely, CFP-1 specifically immunoprecipitated SET-2/SET1 and SWD-2.1/WDR82, but not SET-16/MLL, consistent with it being a unique component of SET1, but not MLL complexes (Clouaire et al. 2014; Lee and Skalnik 2005; Lee et al. 2007). These results define distinct SET-2/SET1 and SET-16/MLL complexes in *C. elegans* embryos (Figure 1C).

WDR-5.1 immunoprecipitates also contained subunits of the NSL histone acetyltransferase (HAT) complex, consistent with findings in other organisms (Dias et al., 2014; Dou et al., 2005; Raja et al., 2010; Zhao et al., 2013). We identified the MOF homologue MYS-2, OGT-1, MCRS-1, and the NSL2 and NSL3 homologues SUMV-1 and SUMV-2, respectively. Homologues of two other NSL components found in other organisms, MBD-R2/PHF20 and NSL1/KANSL1, are not found in the *C. elegans* genome (Hoe and Nicholas 2014). Therefore, as in other species, *C. elegans* WDR-5.1 is found in the NSL complex as well as COMPASS/MLL complexes.

### *C. elegans* homologs of the Sin3S complex copurify with CFP-1 and WDR-5.1

Four additional proteins, SIN-3, HDA-1, MRG-1, and ATHP-1, were reproducibly identified as top hits in both HA::WDR-5.1 and CFP-1::GFP immunoprecipitates (Figure 1B and Table S1). These are homologs of subunits of the Rpd3/Sin3 small complex in yeast (Rpd3/Sin3S) and SHMP in mammalian cells (Figure 1B, see below)(Carrozza et al. 2005; Jelinic et al. 2011). In yeast and other organisms a second type of Rpd3/Sin3 complex is found (Rpd3L in yeast) defined by the presence of SAP30, SDS3, and other subunits (Alland et al., 2002; Fleischer et al., 2003; Lechner et al., 2000; Sardiu et al., 2014; Spain et al., 2010; Wysocka et al., 2003). Because *C. elegans* does not have counterparts of SAP30 and SDS3, it may be that it harbors only a single type of Sin3 complex. We will refer to the complex of SIN-3, HDA-1, MRG-1 and ATHP-1 as the *C. elegans* SIN3 complex (Figure 1C).

Functions ascribed to Rpd3/Sin3 complexes are varied and appear to be context dependent. Although typically referred to as corepressor complexes due to the presence of a histone deacetylase subunit, Rpd3/Sin3 complexes have been associated with both activation and repression of gene expression (Cantor and David, 2017; Cheng et al., 2014; Halder et al., 2017; Lewis et al., 2016; Liu and Pile, 2016; Saha et al., 2016; Saunders et al., 2017; van Oevelen et al., 2008, 2010). In addition, the yeast Rpd3/Sin3S complex has been shown to repress cryptic transcription initiation in transcribed regions and to suppress antisense transcription initiation at promoters (Carrozza et al. 2005; Churchman and Weissman 2011).

Sin3 proteins, which lack known DNA-binding motifs or enzymatic activity, are characterized by the presence of four paired amphipathic helices (PAH) with structural similarity to Myc family transcription factors (Kadamb et al. 2013), and a conserved HDAC-interacting domain (HID) (Laherty et al. 1997). While mammals contain two Sin3 proteins (Sin3A and Sin3B) that share both overlapping and distinct functions (Ayer et al. 1995; Cowley et al. 2005; Dannenberg et al. 2005; David et al. 2008), SIN-3 is the only *C. elegans* homologue. It contains a HID domain, and a single PAH most closely related to the highly conserved PAH1 in mammals (Sahu et al. 2008). *C. elegans* HDA-1 is one of three class I histone deacetylases (HDACs) in *C. elegans* and a component of several other chromatin complexes, as in other species (Kelly and Cowley, 2013; Passannante et al., 2010). MRG-1, the *C. elegans* counterpart of the chromo-domain (CD) protein Eaf3/MRG15, is also found in additional chromatin complexes (Bleuyard et al., 2017; Chen et al., 2010; Huang et al., 2017; Iwamori et al., 2016; Smith et al., 2013), and ATHP-1 (AT Hook plus PHD finger transcription factor), a counterpart of Rco1/Pf1, contains an AT Hook domain that is not found in either Rco1 or Pf1 (Figure S2D).

Western blot analysis on CFP-1::GFP immunoprecipitates using antibodies against endogenous MRG-1, HDA-1, and SIN-3 proteins confirmed the interactions between CFP-1 and SIN3 complex components detected by mass-spectrometry (Figure S2, E and F). We also confirmed that HDA-1 co-precipitates with WDR-5.1 (Figure S2G). We further found that interaction of HDA-1 and MRG-1 with CFP-1 is not dependent on endogenous SIN-3, as both proteins are found in CFP-1 immunoprecipitates obtained from *sin-3* mutant extracts (Figure S2F). We conclude that CFP-1 physically interacts with a Sin3 complex, but may also interact with HDA-1 and MRG-1 in other contexts.

### Subunits of the SET-2/SET1 and SIN3 complex physically interact

We used a yeast two-hybrid assay to assess potential physical interactions between components of the SIN3 and SET-2/SET1 complexes (Fields and Song 1989). A full-length cDNA of each SET-2/SET1 and SIN3 complex subunit was cloned into vectors to express DNA-binding (DB) and activation domain (AD) fusions. Western blot analysis confirmed expression of all cDNAs with the exception of *set-2* (Figure S3A). Testing pairwise interactions of BD and AD fusions by cross-mating, we detected interaction between DPY-30 and ASH-2, and DPY-30 homodimerization within the SET-2/SET1 complex, consistent with studies in other systems (Cho et al., 2007; Dehe et al., 2006; South et al., 2010; van Nuland et al., 2013; Wang et al., 2009b)(Figure 2A). In addition, we detected CFP-1 homodimerisation (Figure 2A). Within the SIN3 complex, we observed an interaction between MRG-1 and ATHP-1, and MRG-1 homodimerization (Figure 2A). Importantly, we found that CFP-1 interacted with the SIN3 complex components ATHP-1 and SIN-3. These results support the above findings that CFP-1 physically interacts with the SIN3 complex.

**Figure 2.**
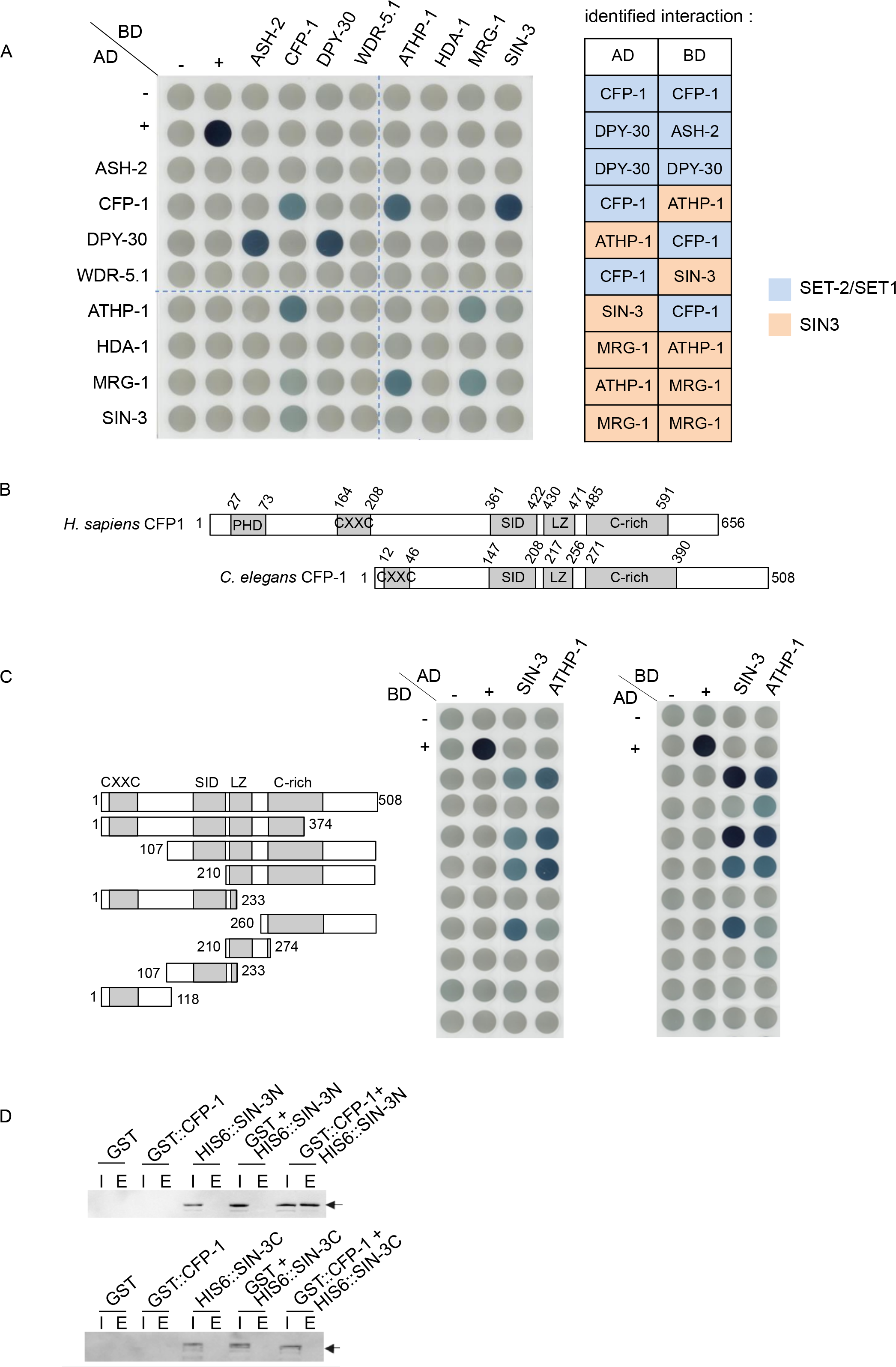
Interaction mapping between subunits of the SET-2/SET1 and SIN3 complexes by Y2H. (A) An interaction matrix was obtained by cross-mating of yeast haploid strains expressing different subunits as AD or BD fusion proteins. Positive control (+) is barnase/barstar interaction (Schreiber and Fersht, 1995). Matings that lead to visually detectable staining in two independent experiments are reported in the tabular form on the right. Most interactions were detected in both directions in the interaction matrix. Failure to detect the DB DPY-30/AD ASH-2 interaction is most likely due to DPY-30 homodimerization in the context of the DB domain fusion interfering with ASH-2 binding. (B) Schematic representation of conserved domains within human and *C. elegans* CFP1 proteins. PHD: plant homeodomain, CXXC: cysteine rich domain, SID: SET1 Interaction Domain, LZ: Leucine Zipper domain predicted by Marcoil software (https://toolkit.tuebingen.mpg.de/#/tools/marcoil). The amino acid position of each domain is denoted above the schematic. (C) Interaction-matrix of full length and truncated CFP-1 tested against full length SIN-3 and ATHP-1. CFP-1 truncations were constructed as BD and AD fusions, as indicated. (D) GST pull-down assay between GST::CFP-1 protein and HIS_6_::SIN-3 N-terminal domain (aa 1-738; left) or HIS_6_::SIN-3 C-terminal domain (aa 699-1507; right). Western blot revealed with anti-Histidine antibodies shows that CFP-1 directly interacts with HIS_6_::SIN3 N-terminal.

### The C-terminal domain of CFP-1 is necessary and sufficient for interaction with SIN-3

Mammalian CFP1 contains an N-terminal PHD domain that recognizes methylated H3K4, a Zn finger CXXC domain that binds to unmethylated CpG dinucleotides, a Set1 interaction domain (SID), a coiled-coiled leucine zipper (LZ) domain, and a cysteine-rich C-terminal domain (Brown et al. 2017; Butler et al. 2008; Mahadevan and Skalnik 2016; Tate et al. 2009; Voo et al. 2000)(Figure 2B and S4). *C. elegans* CFP-1 contains all of these except for the PHD domain. To identify the domains that mediate interaction of CFP-1 with SIN-3 and ATHP-1, we expressed different regions of CFP-1 and tested their ability to interact with full length SIN-3 and ATHP-1 by Y2H as described above. Western blot analysis confirmed the expression of all constructs with the exception of DB 1-374 (Figure S3B). We found that neither the N-terminal CXXC domain, nor the SID domain, were required for interaction with either SIN-3 or ATHP-1 (Figure 2C). The cysteine-rich C-terminal domain fragment interacted with SIN-3, and a larger fragment additionally containing the LZ domain was sufficient for interaction with ATHP-1. These results indicate that CFP-1 binds to SIN-3 through a region containing the cysteine-rich domain, and that interaction with ATHP-1 requires both this region and the LZ domain. We also localized the region of SIN-3 that interacts with CFP-1 to an N-terminal fragment that contains the highly conserved PAH1 domain but lacks the HID domain (HDAC Interaction Domain) (Figure S5).

### Phenotypic similarity of SIN3 and SET-2/SET1 complex mutants

The physical interactions between CFP-1 and SIN3 complex components suggest that they may function in shared processes. To investigate this, we compared phenotypes of *set-2*, *cfp-1*, and *sin-3* mutants alone or in double mutant combinations, using null or strong loss of function alleles for all three genes (Choy et al. 2007; Robert et al. 2014; Xiao et al. 2011). Similar to *set-2* mutants, we observed that *cfp-1* and *sin-3* mutants also have reduced brood size at 20° C (Li and Kelly 2011; Robert et al. 2014; Xiao et al. 2011). Brood size in *cfp-1* mutants showed extreme variability, with some animals showing a near-wild-type brood size, and others producing as few as 10 progeny (Figure 3A). The brood size of *set-2;cfp-1* double mutants is not reduced further compared to *cfp-1* single mutants, suggesting that SET-2 does not have CFP-1 independent fertility functions. However, *sin-3* has fertility functions independent or partially redundant with *set-2* and *cfp-1*, as brood size of *set-2;sin-3* and *cfp-1;sin-3* double mutants is lower than that of the single mutants, with the latter showing fully penetrant sterility (Figure 3A and data not shown). All single and double mutants also have a low level of embryonic lethality (Figure 3B).

**Figure 3.**
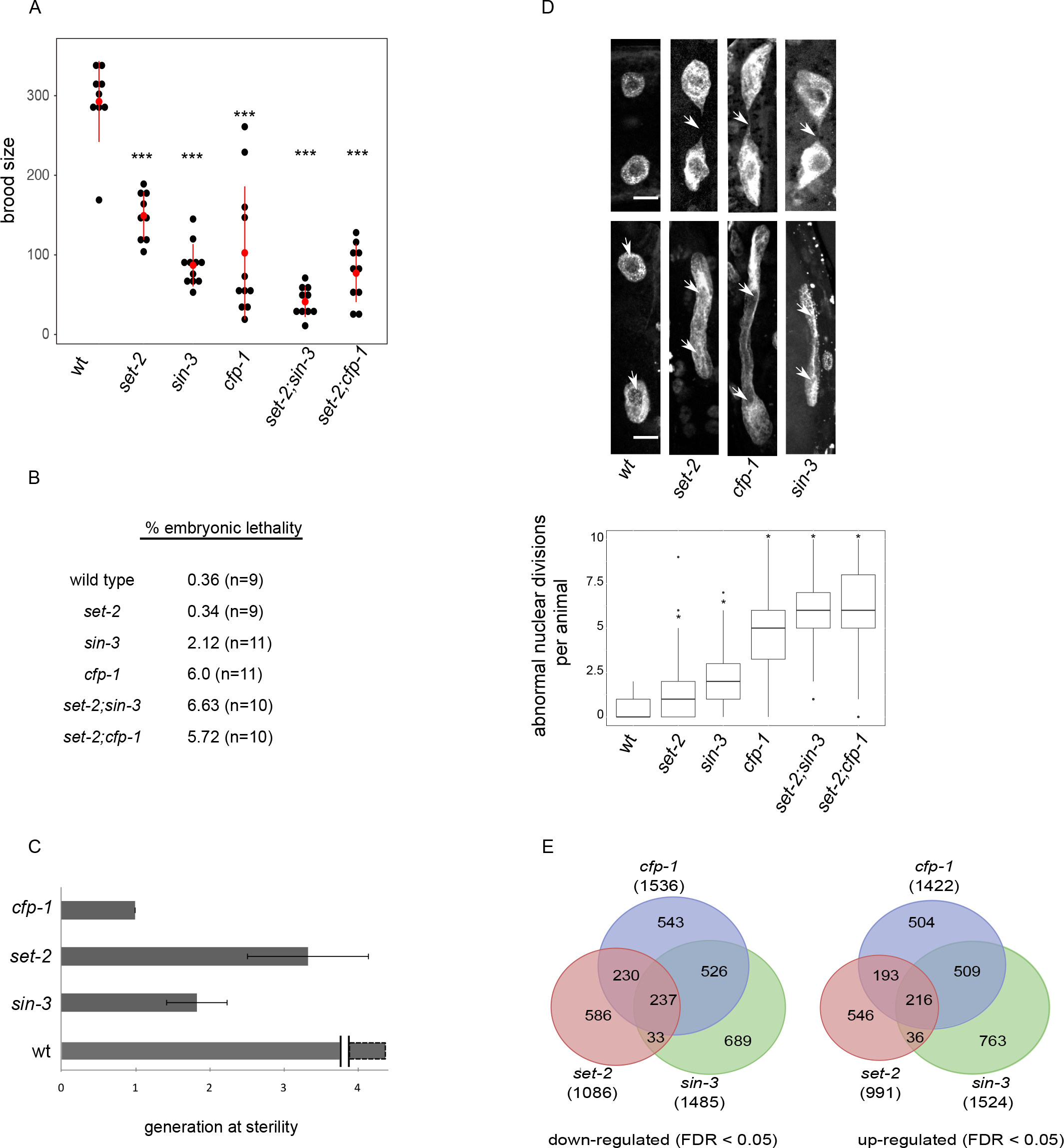
*set-2, cfp-1* and *sin-3* loss of function results in similar phenotypes and changes in gene expression. (A) Total number of progeny of single and double mutant animals of the given genotype. Brood size multiple comparison was done using Dunn-Bonferroni post hoc method following a significant Kruskal Wallis test; asterisks indicate a significant difference from wildtype (*** p ≤ 10e-4). (B) embryonic lethality of single and double mutant strains; n= number of animals scored. (C) Fertility assays of *set-2, cfp-1* and *sin-3* mutants grown at 25°C. Scoring was based on 2 to 3 biological replicates. Wildtype animals can be maintained for more that 40 generations without loss of fertility. (D) Confocal images of DAPI-stained intestinal nuclei from young adults obtained from L1 larva grown at 25°C for 48 hours. Examples of nuclear division abnormalities giving rise to thin chromatin bridges (arrows, top panel), or thick chromatin dense regions connecting two nuclei (arrowheads, bottom panel). Box plots show the total number of segregation defects (thin and thick chromatin bridges) per animal in single and double mutants of the given genotype (n=150 worms for each strain). Multiple comparison was done using Tukey’s ‘Honest Significant Difference’ method following a significant one way analysis of variance test. Asterisks indicate a significant difference from wildtype (* p≤ 5×10e-4). (E) Venn diagram showing the overlap between *cfp-1*, *set-2* and *sin-3* downregulated and upregulated genes.

*set-2* mutants show transgenerational sterility at the stressful temperature of 25° C (Robert et al. 2014), and we found that *cfp-1* and *sin-3* mutants also show this phenotype at 25° C. As expected, *set-2* mutants became sterile at generation F3-F4 (Figure 3C; Robert et al. 2014; Xiao et al. 2011). We observed that *sin-3* mutants become sterile at the F2 generation, whereas the progeny of *cfp-1* mutants that were shifted to 25° C at the L4 stage were sterile (F1 generation).

We also observed that *cfp-1*, *set-2*, and *sin-3* mutants have chromosome segregation defects in intestinal cells that become binucleate in the L1 stage (Hedgecock and White, 1985). Intestinal nuclei were frequently connected by either thin or thick chromatin bridges in single mutants, and often completely failed to separate in *cfp-1* single, and *set-2;sin-3* and *set-2;cfp-1* double mutants (Figure 3D). In summary, the similar phenotypes and genetic interactions suggest that SET-2/SET1 and SIN3 complexes functionally cooperate in the germline and soma.

### Loss of *set-2, cfp-1*, or *sin-3* causes similar effects on gene expression

To ask whether SET-2/SET1 and SIN3 complexes have common roles in gene expression, we next performed RNA-sequencing (RNA-seq) on staged *cfp-1*, *set-2* and *sin-3* mutant embryos. Using DESeq2 (FDR<0.05), we derived lists of differentially expressed genes in each mutant background, finding a similar number that were up- or down-regulated (Table S2 and Figure S6). The three datasets strongly overlap within the up-regulated and down-regulated genes, with 46-65% of differentially expressed genes from each mutant overlapping with those of at least one other mutant, and 14-22% misregulated in all three mutants, consistent with shared regulatory functions (Figure 3E). In addition, pairwise comparisons identified separate sets of genes misregulated only in *cfp-1* and *set-2* mutants, or only in *cfp-1* and *sin-3* mutants (Figure 3E). In contrast, few genes were misregulated only in *set-2* and *sin-3* mutants. Gene ontology (GO) term analysis showed enrichment for biological pathways related to translation, reproduction and embryonic development in all three mutant contexts (Table S3). Downregulation of genes related to reproduction most likely reflects maternally inherited transcripts whose expression is altered in the germline of these mutants (Robert et al. 2014). The patterns of shared gene expression differences in the mutants indicate overlapping roles for SET-2/SET1 and SIN3 complexes, and that CFP-1 also has independent roles with either SIN-3 or SET-2.

### CFP-1 and SET-2/SET1 are needed for H3K4me3 at promoters

Previous studies showed that *cfp-1* or *set-2* inactivation results in greatly reduced global levels of H3K4me3 (Figure S7A, Li and Kelly 2011; Simonet et al. 2007). In addition, CFP-1 binding sites were shown to map to H3K4me3 marked promoters (Chen et al 2014). To determine the roles of the two proteins on the pattern of H3K4me3 at CFP-1 sites, we compared H3K4me3 ChIP-seq signals in wildtype with those in *cfp-1* and *set-2* null mutant embryos, using a spike-in control for normalization (Figure 4A and S7B, see Methods). Using hierarchical clustering, we observed two classes of CFP-1 binding sites in wild-type embryos. Sites with a high level of CFP-1 are strongly marked by H3K4me3, whereas sites with lower CFP-1 levels have low H3K4me3 marking (Figure 4A, B). Based on H3K4me3 levels, we define the high level CFP-1 sites as strong COMPASS targets, and the low level CFP-1 sites as weak COMPASS targets. The finding that the genomic distribution of H3K4me3 is similarly reduced in *cfp-1* and *set-2* mutants confirms that CFP-1 is needed for SET-2 activity at promoters.

**Figure 4.**
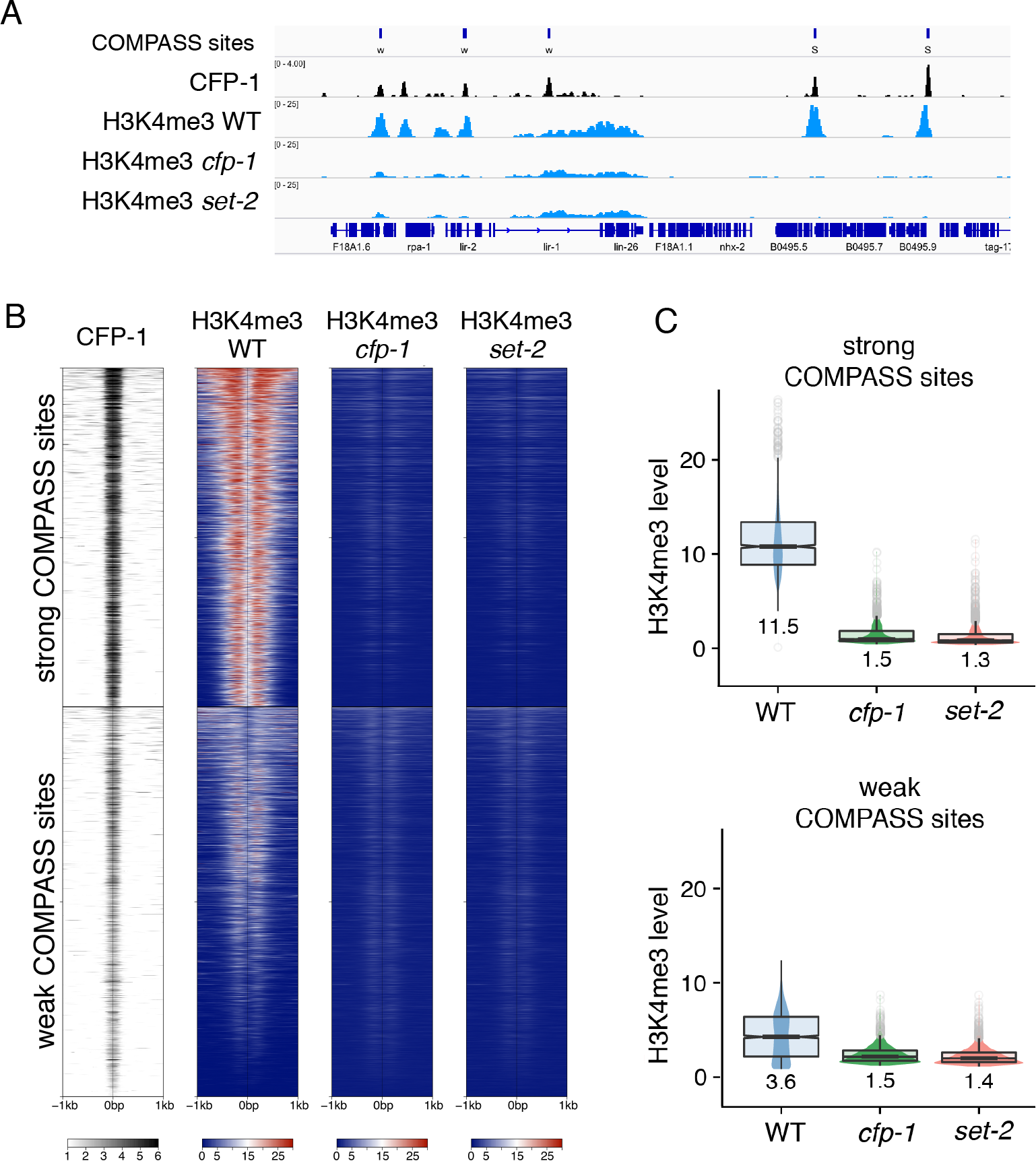
H3K4me3 at strong and weak COMPASS targets is dependent on CFP-1 and SET-2. (A) IGV browser view showing z-score BEADS normalized ChIP-seq signals of CFP-1::GFP, and H3K4me3 in wildtype, *cfp-1*, and *set-2* mutant embryos normalized to *C. briggsae* spike-in (see Methods). Top track shows locations of strong (S) and weak (W) COMPASS targets. (B) Heatmap of CFP-1::GFP, and H3K4me3 in wildtype, *cfp-1*, and *set-2* mutant embryos, using same tracks as in (A). (C) Quantification of H3K4me3 signal (*C. briggsae* normalized) in strong and weak COMPASS targets.

We found that there was no clear relationship between gene expression changes and promoter association of CFP-1. Similar to findings in ES cells (Brown et al 2017), genes downregulated in *cfp-1* and *set-2* mutants were weakly enriched for harboring CFP-1 promoter peaks (Figure S8). However the vast majority of genes with CFP-1 peaks (n=3792) were not significantly altered in expression in any of the three mutants. The lack of a strong association between binding and gene expression regulation suggests that additional factors influence the impact on transcription.

### SIN3 complex components colocalize with CFP-1 at promoter regions

We next investigated how the distribution of SIN3 complex components relates to that of CFP-1. Using ChIP-seq analysis of SIN-3 in wild-type embryos, we observed that the pattern of SIN-3 binding was highly similar to that of CFP-1, with 90% of SIN-3 peaks overlapping a CFP-1 peak (Figures 5A-C). In addition, as observed for CFP-1, SIN-3 levels are higher at strong COMPASS targets than at weak COMPASS targets (Figure 5B). We next determined the distribution of the SIN3 complex component HDA-1. We observed that HDA-1 was also present at most CFP-1 binding sites, with similar levels at strong and weak COMPASS targets (Figure 5A, B). HDA-1 is additionally found at many sites that lack CFP-1 and SIN-3, presumably through its association with other proteins and complexes (Figure 5C)(Passannante et al. 2010).

**Figure 5.**
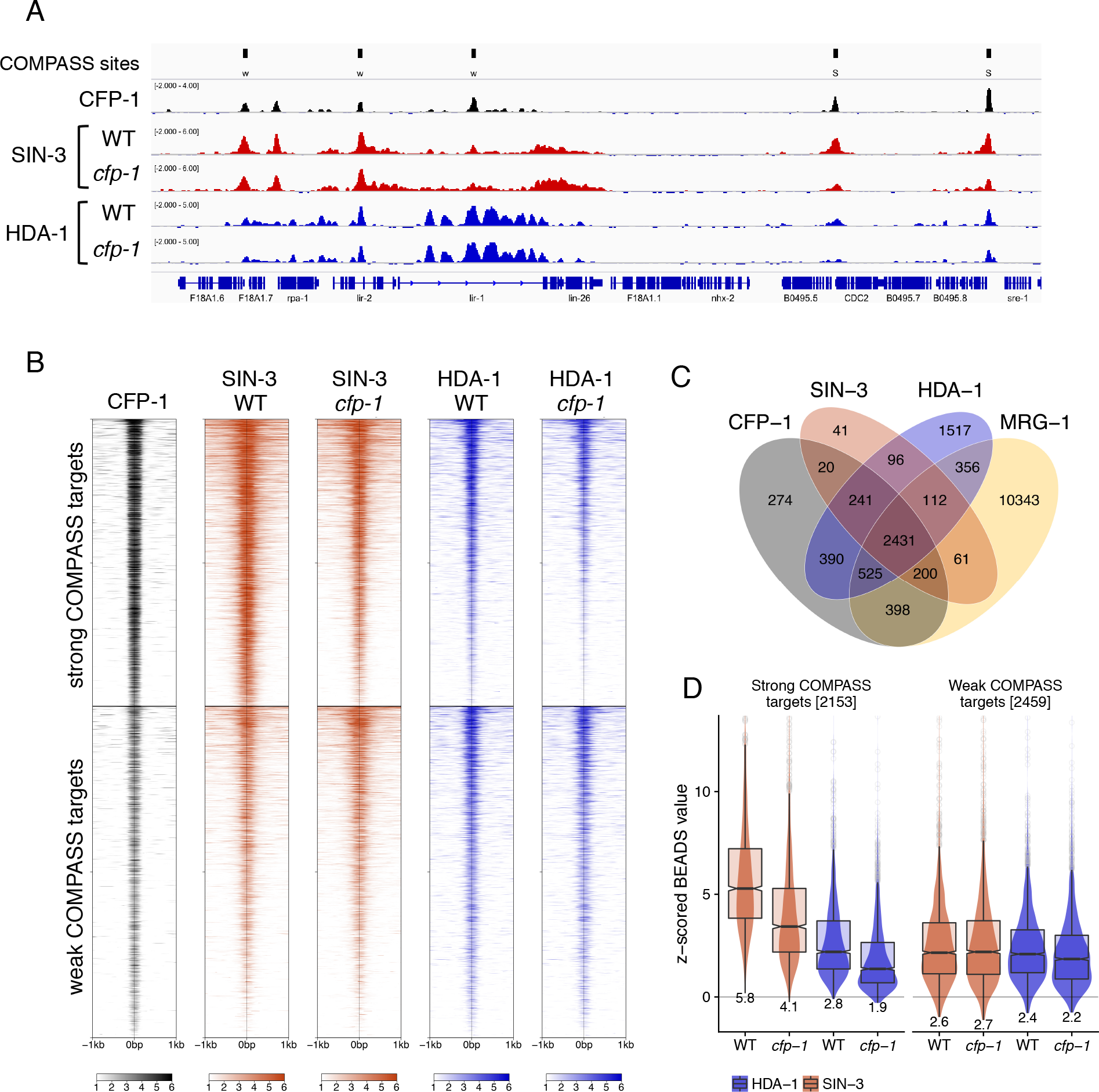
SIN-3 and HDA-1 require CFP-1 for recruitment to strong COMPASS targets. (A) IGV browser view showing z-scored BEADS normalized ChIP-seq signals from mixed embryos of indicated strains. Top track shows locations of strong (S) and weak (W) COMPASS targets. (B) Heatmap of z-scored BEADS normalized ChIP-seq signals from mixed embryos over strong and weak COMPASS targets in indicated strains. (C) Venn diagram showing overlap of CFP-1, SIN-3, HDA-1, and MRG-1 ChIP-seq peaks. (D) Quantification of normalized z-scored SIN-3 and HDA-1 signal in wildtype and *cfp-1* mutants at strong and weak COMPASS targets.

Previously published ChIP-seq data mapping MRG-1 in embryos (Ho et al. 2014) showed weak enrichment at promoters and a broad distribution on the gene bodies of many actively transcribed genes (Figure S9). We found that 91% of sites harboring SIN-3, HDA-1, and CFP-1 peaks (n=2672) contained MRG-1 (Figure 5C). Additionally, we observed that SIN3 complex components SIN-3 and HDA-1 have a broader distribution than CFP-1 and were weakly enriched on gene bodies (Figure S9). The finding that CFP-1 and SIN3 complex components extensively colocalize at promoter regions support connected functions.

### CFP-1 facilitates SIN-3 binding to H3K4me3 enriched promoter regions

The similarity in binding patterns together with our biochemical studies showing that CFP-1 physically associates with the SIN3 complex suggests a potential role in SIN3 chromatin recruitment. To investigate this possibility, we used ChIP-seq to map SIN-3 and HDA-1 binding in *cfp-1* mutant embryos. We observed that strong COMPASS targets had significantly reduced levels of both SIN-3 and HDA-1 in *cfp-1* mutants compared to wildtype (Figures 5A, B, D). In contrast weak COMPASS targets were largely unaffected (Figures 5A, B, D). HDA-1 sites that lack CFP-1 or SIN-3 binding and random genomic regions also showed no change in SIN-3 or HDA-1 levels in *cfp-1* mutants (Figure S10). Together with the physical interaction results, we conclude that CFP-1 plays a direct role in recruiting the SIN3 complex to strong COMPASS target sites.

## Discussion

In this study we identify a physical and functional interaction between CFP-1, the chromatin targeting subunit of the highly conserved SET1/COMPASS complex, and a SIN3 histone deacetylase complex similar to yeast Rpd3S and mouse SHMP containing SIN-3, HDA-1, MRG-1 and ATHP-1. We show that CFP-1 mediates interaction with the SIN3 complex through a direct interaction with SIN-3 and that it promotes recruitment of SIN-3 and HDA-1 at promoter regions. Our results indicate a novel function for CFP-1 in bridging interaction between the SET1/COMPASS H3K4 methyltransferase complex and the SIN3 histone deacetylase complex.

In other organisms, Sin3 proteins are found in multiple complexes of heterogeneous composition (Chaubal and Pile 2018). At least two general types of SIN3 complex exist, which contain SAP30 and SDS3 homologs, or like Rpd3S, Rco1/Pf1 and Eaf3/MRG15 homologs. *C. elegans* may harbor only the Rpd3S type because SAP30 and SDS3 homologs are not present.

Consistent with the presence of multiple H3K4 HMT complexes in metazoans (Shilatifard, 2012), our biochemical data provide evidence for distinct SET-2/SET1 and SET-16/MLL related complexes in *C. elegans*. WDR-5.1 and CFP-1 both immunoprecipited the core complex proteins RBBP-5, ASH-2, and DPY-30, as well as the SET1/COMPASS subunits SET-2/SET1 and SWD-2.1/WDR82 (Dou et al. 2006; Patel et al. 2009, 2011; Shinsky et al. 2015). In addition, WDR-5.1, but not CFP-1, immunoprecipitated unique subunits of the previously identified SET-16/MLL complex including the histone H3K27 demethylase UTX-1, and PIS-1 (Vandamme et al., 2012). WDR-5.1 also co-immunoprecipitated the NSL complex, consistent with its role as a central hub in several additional chromatin-associated complexes (Cai et al. 2010; Cho et al. 2007; Suganuma et al. 2008). Interestingly, in mammalian cells NSL has been shown to promote H3K4me2 activity by MLL1 (Zhao et al., 2013), and we identified the single MLL1 homologue SET-16 with NSL subunits in our experiments, suggesting this activity may be conserved in *C. elegans*.

Y2H and pull-down assays showed a direct interaction between CFP-1 and SIN-3 dependent on the C-terminus of CFP-1 containing the conserved cysteine-rich domain, and the N-terminal domain of SIN-3 containing the highly conserved PAH domain, but not the HDAC-interacting domain (HID). The PAH1 and PAH2 domains of mammalian SIN3 have been shown to facilitate SIN3 recruitment by transcription factors (Sahu et al. 2008). Little is known about the function of the cysteine-rich C-terminal domain of mammalian CFP1. Y2H analysis also confirmed physical interactions within SIN3 and SET-2/SET1 complexes and showed CFP-1 homodimerization, supporting studies suggesting dimerization of CFP1 within the SET1A/B complexes in human cells (van Nuland et al., 2013).

Supporting a functional role of the physical association between SET-2/SET1 and SIN3 complexes, genetic analysis and gene expression studies of *set-2, cfp-1*and *sin-3* mutant animals revealed similar germline and somatic phenotypes, and misregulation of common genes. These findings, together with the physical association and promoter co-occupancy of CFP-1 and SIN-3, suggest that SET-2/SET1 and SIN3 play coordinated roles in modulating gene expression. Previous studies further support common functions for these complexes and other proteins isolated in our proteomics approach. For example, inactivation of the SET-2/SET1 complex subunits *cfp-1, wdr-5.1, dpy-30*, the SIN3 complex subunits *sin-3* and *mrg-1*, and the NSL complex subunits *sumv-1* and *sumv-2* can all suppress the synthetic multivulval (SynMuv) phenotype resulting from mutations in repressive chromatin factors including homologs of the Mi2/NuRD complex, and the Heterochromatin Protein 1 (HP1) homologue HPL-2 (Cui et al. 2006; Yücel et al. 2014). A subset of these genes, including *dpy-30, wdr-5 mrg-1* and *sin-3* also suppress the larval lethality resulting from inactivation of *lin-35*/Rb in a sensitized background (Fay and Yochem 2007).

We found that CFP-1 promotes binding of SIN-3 and HDA-1 at strong COMPASS dependent promoters, consistent with direct recruitment in the context of a SET-2/SET1 complex. SIN-3 binding at weak COMPASS regions was not affected. We note that the SIN-3-interacting domain we identified in the C-terminal of CFP-1 is distinct from the conserved SET1-interacting domain (SID), suggesting that CFP-1 could potentially interact with both SIN-3 and SET-2 at the same time. Interestingly, in mammalian cells the cell proliferation transcription factor Hcf1 was shown to bind the SET1/COMPASS subunit ASH2 and SIN3, supporting a functional connection between the complexes (Wysocka et al. 2003).

A prevailing view is that the regulatory functions of SET1/COMPASS and Rpd3/Sin3 complexes are context dependent, but their mechanisms are not well understood (Chaubal and Pile, 2018; Howe et al., 2017). Given the biochemical activities of complex components, it is plausible that co-occupancy alters gene expression through changes in acetylation and methylation dynamics. In yeast, Set1 dependent methylation was shown to promote deacetylation and repression at promoter regions through the Hst1 HDAC (Kim and Buratowski 2009), while in different species, SIN3 and the Lid/KDM5 H3K4 demethylase copurify and are found together at promoters (Moshkin et al., 2009; Spain et al., 2010; Vermeulen et al., 2010), resulting in H3K4me3 demethylation, deacetylation, and repression of target genes (Moshkin et al. 2009). Additional data suggest that H3K4me3 is able to recruit Sin3 through the ING1 PHD finger protein during myogenesis (Cheng et al. 2014; van Oevelen et al. 2008). Whether H3K4me3 is catalyzed by SET1 or MLL in this context was not investigated.

These and other studies support the view that transcriptional outcome of SET1/COMPASS and Rpd3/Sin3 complexes depends on the nature of other interacting regulators and the chromatin context. Consistent with this, knock-out of Sin3 in different systems results in both gene activation and repression (Gajan et al. 2016; Saha et al. 2016; Saunders et al. 2017; Yao et al. 2017), and we observed no clear relationship between gene expression changes and SIN-3 binding. Similarly, loss of CFP1 or SET1 in different systems causes surprisingly few gene expression changes relative to the number of genes marked by H3K4me3, and no clear relationship is found between expression and marking (Brown et al. 2017; Clouaire et al. 2014; Ramakrishnan et al. 2016; Weiner et al. 2012). In addition to context dependent roles in activating or repressing transcription, it has also been proposed that SET1/COMPASS functions in maintaining transcriptional stability (Howe et al. 2017). Future work on defined loci will be needed to understand these regulatory functions. Because of the high degree of conservation between mammalian and *C. elegans* SET1/COMPASS and Rpd3/Sin3 complexes, our findings that they physically and functionally interact contributes towards understanding the complexity of interactions between chromatin associated proteins with distinct activities.

## Materials and Methods

### Strains and maintenance

Nematode strain maintenance was as described previously (Brenner, 1974). The wild-type strain N2 (Bristol) was used as the reference strain. The strains used are as follows: PFR506 *qaIs22[HA::wdr-5.1; Cbunc-119(+)];wdr-5.1(ok1417)III;* JA1597 *pdpy-30::cfp-1::GFP;* PFR572 *qaIs22[HA::wdr-5.1;Cbunc-119(+)]; wdr-5.1(ok1417)III; pdpy-30::cfp-1::GFP;* PFR625 *qaIs22[HA::wdr-5.1; Cbunc-119(+)], wdr-5.1(ok1417)III; pdpy-30::cfp-1::GFP; sin-3(tm1276)I;* PFR510 *set-2(bn129)/qC1 dpy-19(e1259)glp-1(q339)[qIs26]III*; PFR624 *cfp-1(tm6369) IV/nT1,[unc?(n754),let-?](IV;V);* PFR391 *wdr-5.1(ok1417)III* out crossed twice; PFR590 *sin-3(tm1276)* out-crossed twice; PFR630 *sin-3(tm1276) I/hT2[bli-4(e937) qIs48] I, III;* PFR629 *set-2(bn129)III;sin-3(tm1276)I;* PFR635 *set-2(bn129)III;cfp-1(tm6369)IV/nT1[unc?(n754),let?](IV;V)*; PFR636 *sin-3(tm1276)I;cfp-1(tm6369)IV/nT1[unc?(n754),let?](IV;V)*. ChIP experiments used PFR253 *set-2(bn129)* (outcrossed 10X) and TM6369 *cfp-1(tm6369)* (outcrossed 4X). Fig. S1 shows expression of HA-WDR-5.1 and rescue of *wdr-5.1(ok1417)*. To generate the PFR624 balanced strain, *cfp-1(tm6369)* animals were crossed with wildtype males to generate heterozygote males that were crossed again with [unc] animals segregating from AV112 (*mre-11(ok179),V/nT1,[unc?(n754),let-?](IV;V))*. [unc] animals were selected and screened by PCR for the presence of the *cfp-1(tm6369)* deletion. To generate the PFR630 balanced strain, *sin-3(tm1276)* animals were crossed with wildtype males to generate *sin-3(tm1276)* heterozygote males that were crossed to GFP(+) animals segregating from *[F44E2.7(tm4302)/hT2]*. GFP(+) animals from this last cross were selected and screened by PCR for the presence of the *sin-3(tm1276)* deletion. Yeast two-hybrid strains were: EGY42 (MATa; *trp1, his3, ura3, leu2*); TB50 (MATα; *trp1, his3, ura3, leu2, rme1*). *cfp-1(tm6369)* is a deletion of 254 bp encompassing intron 4, exon 5 and intron 5 of *cfp-1*. It is predicted to produce a truncated CFP-1 protein of 374 aa lacking part of the conserved C-terminal domain. Primers used genotyping are listed in Table S4.

### Immunoprecipitation for proteomics

Immunoprecipitations were performed on frozen embryos prepared by hypochlorite treatment from strains grown at 20°C on enriched NGM. For all immunoprecipitations, wildtype embryos (N2) were treated in parallel to serve as negative control in the mass spectrometry analysis. For HA ::WDR-5.1 immunoprecipitations, embryos from PFR506 were flash-frozen immediately after hypochlorite treatment. For CFP-1::GFP immunoprecipitations, PFR572 late stage embryos were obtained by incubating the embryos collected by hypochlorite treatment for 4h prior to flash freezing in liquid nitrogen. For each condition embryos were ground to powder, resupended in IP buffer (50mM Hepes/KOH pH 7,5; 300 mM KCl; 1mM EDTA; 1 mM MgCl2; 0.2% Igepal-CA630 and 10% glycerol) containing complete protease inhibitors [Roche] and 1 mM PMSF, and sonicated. Protein extracts were recovered in supernantant following centrifugation at 20 000 g for 15 min at 4°C an 20°C and flash frozen in liquid nitrogen. Protein concentrations were estimated using the Bradford assay [BIO-RAD Protein Assay Dye]. For HA::WDR-5.1 immunoprecipitation, approximately 60 mg of total protein extract was incubated with protein G agarose beads [Sigma-Aldrich] in Bio-Spin Chromatography columns [BioRAD] for 30 min at 4°C on a rotator. Flow-through was collected and incubated with 240 *μ*l slurry of anti-HA affinity matrix beads [Roche] in a fresh Bio-Spin Chromatography column for 90 min at 4°C on a rotator. The matrix was washed three times in IP buffer at 4°C and once in Benzo buffer (Hepes/KOH 50mM pH 7,5; KCl 150mM; EDTA 1mM; MgCl2 1mM; Igepal-CA630 0.2%; glycerol 10%). The matrix was then incubated in 400 *μ*l of Benzo buffer containing 2500 units of benzonase [Sigma] for 1 h at 4°C and washed three times in IP buffer. Four successive elutions were performed at 37°C for 15 min each with HA peptide (250 *μ*g/ml in 240 *μ*l of IP buffer). The first three eluates were pooled and concentrated 20 times (final volume 35 *μ*l) using Amicon Ultra centrifugal device [Merck]. 1/70 and 1/700 of this eluate were resolved on a 4–12% NuPage Novex gel [Thermo Fischer] and the gel either stained with SilverQuest staining kit [Thermo Fischer] or analyzed by western blot with anti-HA antibody [Covance HA.11, clone 16B12]. 33 *μ*l of the eluate was diluted with 11 *μ*l of LDS4X buffer [Thermo Fischer] and analyzed by mass spectrometry. For CFP-1::GFP immunoprecipitation, approximatively 70 mg of total protein were incubated in IP buffer with 100 *μ*l of GFP-TRAP MA beads slurry [Chromotek] for 3h at 4°C on a rotator. Beads were collected with a magnet, washed three times in IP buffer and one time in Benzo buffer, and then treated with benzonase. Eluates were recovered by incubation at 95°C for 10 min in 60 *μ*l of LDS 1X buffer. 1/10 and 1/50 of this eluate were resolved on a 4–12% NuPage Novex gel [Thermo Fischer] and either stained with SilverQuest staining kit [Thermo Fischer] or analyzed by western blot with anti-GFP antibody [Sigma, 11814460001, clones 7.1 and 13.1] respectively, and 40 *μ*l of the eluate was analyzed by Mass spectrometry.

### Mass spectrometry-based proteomic analyses

Proteins were stacked in the top of a SDS-PAGE gel (4-12% NuPAGE, Life Technologies) and stained with Coomassie blue R-250 before in-gel digestion using modified trypsin (Promega, sequencing grade) as previously described (Casabona et al., 2013). Resulting peptides were analyzed by online nanoLC-MS/MS (UltiMate 3000 and LTQ-Orbitrap Velos Pro, Thermo Scientific). For this, peptides were sampled on a 300 *μ*m x 5 mm PepMap C18 precolumn and separated on a 75 *μ*m x 250 mm C18 column (PepMap, Thermo Scientific). MS and MS/MS data were acquired using Xcalibur (Thermo Scientific). Peptides and proteins were identified using Mascot (version 2.5.1) through concomitant searches against Uniprot (*Caenorhabditis elegans* taxonomy), classical contaminants database (homemade) and the corresponding reversed databases. The Proline software (http://proline.profiproteomics.fr) was used to filter the results (conservation of rank 1 peptides, peptide identification FDR < 1% as calculated on peptide-spectrum match scores by employing the reverse database strategy, minimum peptide score of 25, and minimum of 1 specific peptide per identified protein group) before performing a compilation, grouping and comparison of the protein groups from the different samples.

### Co-immunoprecipitation experiments

Co-immunoprecipitations with CFP-1::GFP were performed starting from 4 mg total protein embryonic extract from the strain containing the two transgenes CFP-1::GFP and HA::WDR-5.1. Samples were processed as in proteomic experiments. Co-immunoprecipitations with HA::WDR-5.1 were performed with the eluates sent to mass spectrometry analysis. Samples were processed as in proteomic experiments. Eluates were boiled in LDS sample buffer and analyzed on 4–12% NuPage Novex gels [Thermo Fischer] or Mini-PROTEAN TGX Stain-Free Precast gels [BIO-RAD] followed by western blotting. Antibodies used were : anti-GFP [Sigma, 11814460001, clones 7.1 and 13.1] (1/1000); anti-HA [Covance HA.11, clone 16B12] (1/2000); anti-HDA-1 [Novus Biologicals, 38660002] (1/2000); anti-DPY-30 [Novus Biologicals, 45110002] (1/5000); anti-ASH-2 (gift from B. Meyer) (1/4000); anti-MRG-1 antibody [Novus Biologicals, 35530002] (1/3000); anti-SIN-3 Q60131017(1/1000).

### Plasmids construction for Y2H

Plasmids used for expression of BD and AD fusions were derived from pEG202 (Clontech; Genbank Accession Number U89960) and pJG4–5 plasmids (Clontech; Genbank Accession Number <u>U89961</u>), respectively (Golemis and Brent, 1992). Constructions were generated by cloning the cDNA of the gene of interest in the *Xho*I restriction site of the pEG202 and pJG4– 5 plasmids using the Gibson method (Gibson, 2011). CFP-1 truncations were obtained by the same reaction using the *cfp-1* cDNA sequence as template. PCR reactions were carried out using pHusion polymerase and primers listed in Table S3. All products were verified by sequencing. pSH18–34, bearing a beta-galactosidase gene under the control of four overlapping LexA operators was used as reporter vector (Estojak et al., 1995).

### Interaction Trap/Two-Hybrid system to identify interacting protein

Y2H assay is based on the LexA (BD)/B42 (AD) system (Golemis and Brent, 1992; Finley and Brent, 1994). Cross-matings were performed in liquid phase (Ito et al., 2001). Competent haploid EGY42a cells were co-transformed with 1*μ*g of pSH18-34 (reporter vector) and 1*μ*g of BD construct. Competent TB50α cells were transformed with 1*μ*g of AD construct. Yeasts were selected for 3 days at 30°C on SD-UH (BD strains) and SD-W (AD strains) medium. Matings were performed overnight at 30°C in liquid YPAD (Kolonin et al., 2000). Cross-mating ensured that each hetero-interaction was tested twice (in both directions of the interaction matrix) and allowed the detection of homodimerisations. Diploids were amplified in selective liquid SD-UHW medium. For β-Galactosidase assays, 50*μ*l of each diploids culture was inoculated (at OD595nm = 6) in 1ml of pre-warmed (25°C) SGR-UHW medium supplemented with X-Gal (Thermo Scientific, #R0404) in Deepwell 96 well plates. Cultures were then incubated for 48h at 25°C, centrifuged 5 min at 192g, resuspended in 300*μ*l, and transferred in flat bottom *μ*Clear Cellstar^®^ plates (Greiner Bio one) for scanning and phenotype assessment.

### Protein expression and GST pull-down assay

Full-length GST::CFP-1 and HIS_6_::SIN-3 fragments (1-738, 699-1507) were aplified using primers listed in Table S3 and subcloned into pGEX-6-P1 [Sigma GE28-9546-48] and pPROEX HTa (gift from Dr L. Terradot, MMSB, Lyon), respectively, using the Gibson method (Gibson 2011). All proteins were expressed in BL21 Rosetta 2 [Merck-Millipore 71402]. Bacteria were grown to OD600 0,6 and protein expression induced with 1mM IPTG at 16°C overnight. The pellet from 1 L of bacterial culture was resuspended in 10 ml Lysis buffer (50mM Tris pH 8,0 ; 300mM NaCl ; 0,1mM EDTA ; 0,1% Triton X-100 ; 0,05% NP-40; 1mM MgCl2; 5% Glycerol) containing protease inhibitor [Roche 05056489001]. Samples were sonicated on ice and centrifuged at 20 000 g for 20 min. For pulldown assays 200uL of GST::CFP-1 supernatant was mixed with 800 uL HIS_6_::SIN-3[1-738] or HIS_6_::SIN-3[699-1507], respectively, and incubated overnight at 4°C on a rotating wheel. Samples were submitted to GST purification on a Biosprint 15 automat from Qiagen. Samples were washed 3 times with Lysis buffer and eluted with 50 uL of Lysis buffer containing 20mM Glutathion. Eluted fractions were analyzed by western blot using a mouse anti-Histidine antibody [Sigma H1029] (1/3000) and Stain-Free gels (Bio-Rad).

### Brood size and embryonic lethality assays

For each strain, 10 L4 worms were isolated to single plates in the presence of excess food at 20°C, and allowed to develop into egg-laying adults overnight. Adult animals were then transferred to fresh plates every 12 hours until they ceased laying eggs. Plates were scored for number of viable progeny and dead embryos that failed to hatch 24 hrs after removal of the mother.

### Fertility assay

Six independent lines were established from freshly thawed *sin-3(tm1276)* animals maintained as homozyotes, and homozygous *set-2(bn129)* and *cfp-1(tm6369)* animals obtained from balanced strains PFR 510 and PFR 624, respectively. For each line, six homozygous L4 stage animals were transferred to single plates with fresh *Escherichia coli*, in the presence of excess food and cultivated at 25°C. From each generation, six worms were again picked to single plates until animals became sterile (fewer than 10 progeny/plate).

### Characterization of nuclear divisions in intestinal nuclei

Adult animals were treated with hypochlorite solution to obtained L1 synchronized larva. L1 larva were transferred to 25°C for 48 hrs, until they developed into adults. Young adults were stained with DAPI staining and analyzed with a Zeiss 710 Confocal Microscope. Experiments were performed in three independent replicates and intestinal nuclei from a total of 150 worms for each strain were scored.

### RNA-seq analysis

RNAs were extracted from frozen early stage embryos prepared by hypochlorite treatment of young adults. Two to three independent biological replicates were performed for each strain. RNAs were extracted with NucleoZol [Macherey-Nagel] according to manufacturer’s instructions and treated with DNAse [Turbo-free DNAse, Ambion]. Integrity of RNA was assessed on Tape Station 4200 [Agilent]. RNA-seq librairies were generated at the GenomEast Platform [IGBMC, Strasbourg, France] using the directional mRNA-Seq SamplePrep [Illumina] and sequenced using the Illumina Hiseq 4000 technology. All RNA-seq data were mapped to the *C. elegans* reference genome (WS254) by RNA-STAR (Version 2.4.1d). Reads below a mapping score of 10 were filtered using SAMtools (Version 0.1.19). Of the 46,771 annotated genes, 20,183 were selected as protein coding genes and among them, 11,630 had sufficient read representation (baseMean > 10) for further analysis. The gene expression level in each sample was calculated by htseq-count (Version 0.7.2) and differential expression between the different strains was calculated with DESeq2 (Version 1.10.1 using R version 3.2.4). Gene expression data are available at GEO with the accession GSE110072. Reviewers can access the data at https://www.ncbi.nlm.nih.gov/geo/query/acc.cgi?acc=GSE110072 using password yjihwieqhzsnnkr.

### Western blot analysis on histone marks

Embryos were obtained by hypochlorite treatment of adults grown on solid media at 20°C and frozen in liquid nitrogen. Pellets were resuspended in buffer (50mM Hepes/KOH pH 7,5; 300mM KCl; 1mM EDTA; 1mM MgCl2; 0.2% Igepal-CA630 and 10% glycerol) containing complete protease inhibitors [Roche] and PMSF (1mM), and sonicated. Total protein amount was quantified by the Bradford assay [BIORAD]. Dilutions of wild-type total protein extracts were analyzed to determine the upper limit of linearity of the following antibodies: anti-H3K4me3 [Diagenode C15310003] (1/2000) and anti-H3 [Active Motif, 39163] (1/20000). Two dilutions of total protein extracts were analyzed by western blot for each strain and each antibody.

### Chromatin immunoprecipitation

Wildtype, *cfp-1(tm6369)*, and *set-2(bn129)* mixed embryos were obtained by growing strains at 20°C in liquid culture using standard S-basal medium with HB101 bacteria. Strains were grown to the adult stage then bleached to obtain embryos, which were washed in M9, then frozen into "popcorn" by dripping embryo slurry into liquid nitrogen. Chromatin immunoprecipitations and library preparations were conducted as in (McMurchy et al. 2017), using formaldehyde as a fixative for the H3K4me3 ChIPs (30ug DNA, 2.5ug antibody) and formaldehyde and EGS as fixatives for the SIN-3 (15ug DNA, 2.5ug antibody) and HDA-1 (30ug DNA, 2.5ug antibody) ChIPs. Approximately 10% *C. briggsae* chromatin extract was spiked into the *C. elegans* extract for the H3K4me3 ChIPs and 5% into the HDA-1 ChIPs. The HDA-1 antibody did not detect *C. briggsae* HDA-1 and so was not used for normalization. Two different antibodies to SIN-3 were raised through Strategic Diagnostics International by DNA immunization using aa427-576 (Q5986 and Q6013). Chromatin immunoprecipitations were conducted in duplicate with both SIN-3 antibodies in wild-type embryos; ChIP-seq patterns using these two SIN-3 antibodies were highly concordant (Figure S11). Comparison of SIN-3 ChIP levels between wild-type and *cfp-1* mutant embryos were done using SIN-3 antibody Q5986. HDA-1 ChIPs were done using Novus 38660002/Q2354 and H3K4me3 ChIPs used Abcam ab8580. The age distributions of mixed embryo collections were in the following proportions (% <300 cell / % over300cell, average of the two replicates): H3K4me3 ChIPs: WT N2, 51/49; *cfp-1*, 59/51, *set-2*, 54/46. SIN-3 and HDA-1 ChIPs: WT N2, 48/52; *cfp-1*, 49/51. Sequencing libraries were constructed as in (McMurchy et al. 2017). ChIP-seq libraries were sequenced using an Illumina HiSeq1500.

### SIN-3, HDA-1, and CFP-1::GFP ChIP-seq data processing

CFP-1::GFP (GEO GSE49870), SIN-3, and HDA-1 ChIP-seq reads were aligned to the ce11 assembly of the *C. elegans* genome using BWA v. 0.7.7 (Li and Durbin 2010) with default settings (BWA-backtrack algorithm). The SAMtools v. 0.1.19 ‘view’ utility was used to convert the alignments to BAM format. Normalized ChIP-seq coverage tracks was generated using the BEADS algorithm (Cheung et al., 2011; Stempor 2014). ChIP-seq peaks were called for SIN-3, HDA-1, and CFP-1::GFP in wild-type embryos using MACS2 v. 2.1.1 (Feng et al. 2012) with a q-value cut-off of 0.05 and fragment size of 150bp against summed ChIP-seq input (GEO GSE87524). Peak summits were extended 150bp upstream and downstream, creating 300bp peak regions. Peaks obtained from each replicate were combined by intersection and extending the resulting regions to 300bp. Peaks overlapping non-mappable (GEM-mappability < 25%; (Derrien et al. 2012) or blacklisted regions (https://gist.githubusercontent.com/Przemol/ef62ac7ed41d3a84ad6c478132417770/raw/56e98b99e6188c8fb3dfb806ff6f382fe91c27fb/CombinedBlacklists.bed) were discarded. SIN-3 peak calls are the intersection of peaks obtained using the Q5986 and Q6013 antibodies. SeqPlots (Stempor and Ahringer 2016) software was used to to separate CFP-1::GFP peaks into strong and weak COMPASS sites (k-means clustering), and to visualise CFP-1::GFP, HDA-1 SIN-3, MRG-1 (GEO GSE50333) and H3K4me3 ChIP-seq tracks as heatmaps. The IGV Genome Browser (Thorvaldsdóttir et al. 2013) was applied to visualise example regions. Strong and weak COMPASS peaks were assigned to genes based on overlap with promoter annotations in (Janes et al 2018). ChIP-seq data generated in this study is available at GSE114715. Reviewers can access the data at https://www.ncbi.nlm.nih.gov/geo/query/acc.cgi?acc=GSE114715 using password ibepaiumjbonhaz.

### Spike-in normalization of H3K4me3 ChIP-seq

Sequencing reads from H3K4me3 ChIP and corresponding input samples were mapped to a concatenated reference genome sequence containing *C. elegans* ce11 and *C. briggsae* cb3 using BWA (Li and Durbin 2010) and were then separated by species. Only reads that mapped uniquely (mapq>=10) to non-blacklisted regions were kept. The spike-in ratios of *C. briggsae* to *C. elegans* chromatin present in the combined extract were calculated from the input sequence as *C. briggsae* read count divided by *C. elegans* read count. *C briggsae* H3K4me3 peaks were called from ChIP data using MACS2 (Feng et al. 2012) with default parameters. Scaling factors for each ChIP samples were calculated as corresponding spike-in ratio divided by *C. briggsae* H3K4me3 ChIP read count in peak regions in millions. These scaling factors were applied to *C. elegans* H3K4me3 ChIP raw coverage track. As a last step, ChIP background was removed from the scale coverage tracks by subtracting the mode and setting negative values to zero. The resulting tracks were used for visualization and analysing H3K4me3 levels.

### ChIP-seq signal quantifications

To compare SIN-3 and HDA-1 binding between wildtype and *cfp-1* mutant embryos, we quantified average BEADS normalised, z-scored signal on different peak sets. The signal was obtained using the *bigWigSummary* utility from Kent library (Kent et al. 2010) implemented in *rtracklayer* package in R. These signals were represented as overlaid violin plots (showing signal distribution) and Tukey box plots (showing estimation of statistical significance of difference between medians as notches) (Turkey, 1977). The comparison of H3K4me3 levels in wt, *cfp-1*, and *set-2* mutants was done in the same way, using spike-in normalized signal tracks for quantification.

## Acknowledgements

We are grateful to G. Benoit for help in bioinformatics analysis, JL. Bessereau and J. Govin for scientific discussion, and R. Margueron for critical reading of the manuscript. We acknowledge the discovery platform and informatics group at EDyP, the PSMN (Pôle Scientifique de Modélisation Numérique) of the ENS de Lyon, and the SFR Biosciences (PLATIM and PSF). Thanks to B. Meyer for ASH2 antibodies and the *Caenorhabditis* Genetic Center, which is supported by the National Center for Research, for strains. Proteomic experiments were partly supported by the Proteomics French Infrastructure (ANR-10-INBS-08-01 grant) and the Labex GRAL (ANR-10-LABX-49-01). F.P. acknowledges support by the ANR (N° 15-CE12-0018-01), the CNRS, and the Fondation ARC. J.A. acknowledges support by a Wellcome Trust Senior Research Fellowship (101863) and core support from the Wellcome Trust (092096) and Cancer Research UK (C6946/A14492).

## Author contributions

F.B. carried out and interpreted biochemistry experiments and genetic analysis, M.C. designed and interpreted biochemistry experiments, and assisted in their execution, C.B. carried out and interpreted expression profiling experiments and analysis, M.H. assisted in genetic analysis, D.C and M.S. provided expertise in Y2H assays, H.P. assisted in RNA-seq analysis, Y.C. carried out mass spectrometry analyses, A.A. carried out ChIP-seq experiments, Y.D. and R.C. collected and validated samples, P.S. analyzed ChIP-seq data, N.H. performed spike-in normalization, F.B., C.B., P.S. made figures, F.P. and J.A. designed experiments, interpreted results and wrote the paper.

## Declaration of Interests

The authors declare no competing interests.

## References

Alland L, David G, Shen-Li H, Potes J, Muhle R, Lee H-C, Hou H, Chen K, DePinho RA. 2002. Identification of mammalian Sds3 as an integral component of the Sin3/histone deacetylase corepressor complex. Mol Cell Biol 22: 2743–2750.

Ardehali MB, Mei A, Zobeck KL, Caron M, Lis JT, Kusch T. 2011. Drosophila Set1 is the major histone H3 lysine 4 trimethyltransferase with role in transcription. EMBO J 30: 2817–2828.

Ayer DE, Lawrence QA, Eisenman RN. 1995. Mad-Max transcriptional repression is mediated by ternary complex formation with mammalian homologs of yeast repressor Sin3. Cell 80: 767–776.

Bernstein BE, Kamal M, Lindblad-Toh K, Bekiranov S, Bailey DK, Huebert DJ, McMahon S, Karlsson EK, Kulbokas EJ, Gingeras TR, et al. 2005. Genomic maps and comparative analysis of histone modifications in human and mouse. Cell 120: 169–181.

Bledau AS, Schmidt K, Neumann K, Hill U, Ciotta G, Gupta A, Torres DC, Fu J, Kranz A, Stewart AF, et al. 2014. The H3K4 methyltransferase Setd1a is first required at the epiblast stage, whereas Setd1b becomes essential after gastrulation. Dev Camb Engl 141: 1022–1035.

Bleuyard J-Y, Fournier M, Nakato R, Couturier AM, Katou Y, Ralf C, Hester SS, Dominguez D, Rhodes D, Humphrey TC, et al. 2017. MRG15-mediated tethering of PALB2 to unperturbed chromatin protects active genes from genotoxic stress. Proc Natl Acad Sci U S A 114: 7671–7676.

Brown DA, Di Cerbo V, Feldmann A, Ahn J, Ito S, Blackledge NP, Nakayama M, McClellan M, Dimitrova E, Turberfield AH, et al. 2017. The SET1 Complex Selects Actively Transcribed Target Genes via Multivalent Interaction with CpG Island Chromatin. Cell Rep 20: 2313–2327.

Butler JS, Lee J-H, Skalnik DG. 2008. CFP1 interacts with DNMT1 independently of association with the Setd1 Histone H3K4 methyltransferase complexes. DNA Cell Biol 27: 533–543.

Cai Y, Jin J, Swanson SK, Cole MD, Choi SH, Florens L, Washburn MP, Conaway JW, Conaway RC. 2010. Subunit composition and substrate specificity of a MOF-containing histone acetyltransferase distinct from the male-specific lethal (MSL) complex. J Biol Chem 285: 4268–4272.

Cantor DJ, David G. 2017. The chromatin-associated Sin3B protein is required for hematopoietic stem cell functions in mice. Blood 129: 60–70.

Carrozza MJ, Li B, Florens L, Suganuma T, Swanson SK, Lee KK, Shia W-J, Anderson S, Yates J, Washburn MP, et al. 2005. Histone H3 methylation by Set2 directs deacetylation of coding regions by Rpd3S to suppress spurious intragenic transcription. Cell 123: 581–592.

Chaubal A, Pile LA. 2018. Same agent, different messages: insight into transcriptional regulation by SIN3 isoforms. Epigenetics Chromatin 11: 17.

Chen M, Tominaga K, Pereira-Smith OM. 2010. Emerging role of the MORF/MRG gene family in various biological processes, including aging. Ann N Y Acad Sci 1197: 134–141.

Chen RA-J, Stempor P, Down TA, Zeiser E, Feuer SK, Ahringer J. 2014. Extreme HOT regions are CpG-dense promoters in C. elegans and humans. Genome Res 24: 1138–1146.

Cheng J, Blum R, Bowman C, Hu D, Shilatifard A, Shen S, Dynlacht BD. 2014. A Role for H3K4 Monomethylation in Gene Repression and Partitioning of Chromatin Readers. Mol Cell 53: 979–992.

Cheung M-S, Down TA, Latorre I, Ahringer J. 2011. Systematic bias in high-throughput sequencing data and its correction by BEADS. Nucleic Acids Res 39: e103.

Cho Y-W, Hong T, Hong S, Guo H, Yu H, Kim D, Guszczynski T, Dressler GR, Copeland TD, Kalkum M, et al. 2007. PTIP associates with MLL3- and MLL4-containing histone H3 lysine 4 methyltransferase complex. J Biol Chem 282: 20395–20406.

Choy SW, Wong YM, Ho SH, Chow KL. 2007. C. elegans SIN-3 and its associated HDAC corepressor complex act as mediators of male sensory ray development. Biochem Biophys Res Commun 358: 802–807.

Churchman LS, Weissman JS. 2011. Nascent transcript sequencing visualizes transcription at nucleotide resolution. Nature 469: 368–373.

Clouaire T, Webb S, Bird A. 2014. Cfp1 is required for gene expression-dependent H3K4 trimethylation and H3K9 acetylation in embryonic stem cells. Genome Biol 15: 451.

Clouaire T, Webb S, Skene P, Illingworth R, Kerr A, Andrews R, Lee J-H, Skalnik D, Bird A. 2012. Cfp1 integrates both CpG content and gene activity for accurate H3K4me3 deposition in embryonic stem cells. Genes Dev 26: 1714–1728.

Cosgrove MS, Patel A. Mixed lineage leukemia: a structure-function perspective of the MLL1 protein. Febs J 277: 1832–42.

Cowley SM, Iritani BM, Mendrysa SM, Xu T, Cheng PF, Yada J, Liggitt HD, Eisenman RN. 2005. The mSin3A chromatin-modifying complex is essential for embryogenesis and T-cell development. Mol Cell Biol 25: 6990–7004.

Cui M, Kim EB, Han M. 2006. Diverse chromatin remodeling genes antagonize the Rb-involved SynMuv pathways in C. elegans. PLoS Genet 2: e74.

Dannenberg J-H, David G, Zhong S, van der Torre J, Wong WH, Depinho RA. 2005. mSin3A corepressor regulates diverse transcriptional networks governing normal and neoplastic growth and survival. Genes Dev 19: 1581–1595.

David G, Grandinetti KB, Finnerty PM, Simpson N, Chu GC, Depinho RA. 2008. Specific requirement of the chromatin modifier mSin3B in cell cycle exit and cellular differentiation. Proc Natl Acad Sci U S A 105: 4168–4172.

Dehe P-M, Dichtl B, Schaft D, Roguev A, Pamblanco M, Lebrun R, Rodriguez-Gil A, Mkandawire M, Landsberg K, Shevchenko A, et al. 2006a. Protein Interactions within the Set1 Complex and Their Roles in the Regulation of Histone 3 Lysine 4 Methylation. J Biol Chem 281: 35404–35412.

Dehe PM, Dichtl B, Schaft D, Roguev A, Pamblanco M, Lebrun R, Rodriguez-Gil A, Mkandawire M, Landsberg K, Shevchenko A, et al. 2006b. Protein interactions within the Set1 complex and their roles in the regulation of histone 3 lysine 4 methylation. J Biol Chem 281: 35404–12.

Denissov S, Hofemeister H, Marks H, Kranz A, Ciotta G, Singh S, Anastassiadis K, Stunnenberg HG, Stewart AF. 2014. Mll2 is required for H3K4 trimethylation on bivalent promoters in embryonic stem cells, whereas Mll1 is redundant. Dev Camb Engl 141: 526–537.

Derrien T, Estellé J, Marco Sola S, Knowles DG, Raineri E, Guigó R, Ribeca P. 2012. Fast Computation and Applications of Genome Mappability ed. C.A. Ouzounis. PLoS ONE 7: e30377.

Dias J, Van Nguyen N, Georgiev P, Gaub A, Brettschneider J, Cusack S, Kadlec J, Akhtar A. 2014a. Structural analysis of the KANSL1/WDR5/KANSL2 complex reveals that WDR5 is required for efficient assembly and chromatin targeting of the NSL complex. Genes Dev 28: 929–942.

Dias J, Van Nguyen N, Georgiev P, Gaub A, Brettschneider J, Cusack S, Kadlec J, Akhtar A. 2014b. Structural analysis of the KANSL1/WDR5/KANSL2 complex reveals that WDR5 is required for efficient assembly and chromatin targeting of the NSL complex. Genes Dev 28: 929–942.

Dou Y, Milne TA, Ruthenburg AJ, Lee S, Lee JW, Verdine GL, Allis CD, Roeder RG. 2006. Regulation of MLL1 H3K4 methyltransferase activity by its core components. Nat Struct Mol Biol 13: 713–719.

Dou Y, Milne TA, Tackett AJ, Smith ER, Fukuda A, Wysocka J, Allis CD, Chait BT, Hess JL, Roeder RG. 2005. Physical association and coordinate function of the H3 K4 methyltransferase MLL1 and the H4 K16 acetyltransferase MOF. Cell 121: 873–85.

Fay DS, Yochem J. 2007. The SynMuv genes of Caenorhabditis elegans in vulval development and beyond. Dev Biol 306: 1–9.

Feng J, Liu T, Qin B, Zhang Y, Liu XS. 2012. Identifying ChIP-seq enrichment using MACS. Nat Protoc 7: 1728–1740.

Fields S, Song O. 1989. A novel genetic system to detect protein-protein interactions. Nature 340: 245–246.

Fleischer TC, Yun UJ, Ayer DE. 2003. Identification and characterization of three new components of the mSin3A corepressor complex. Mol Cell Biol 23: 3456–3467.

Gajan A, Barnes VL, Liu M, Saha N, Pile LA. 2016. The histone demethylase dKDM5/LID interacts with the SIN3 histone deacetylase complex and shares functional similarities with SIN3. Epigenetics Chromatin 9: 4.

Gibson DG. 2011. Enzymatic Assembly of Overlapping DNA Fragments. In Methods in Enzymology, Vol. 498 of, pp. 349–361, Elsevier http://linkinghub.elsevier.com/retrieve/pii/B9780123851208000152 (Accessed July 11, 2017).

Glaser S, Schaft J, Lubitz S, Vintersten K, van der Hoeven F, Tufteland KR, Aasland R, Anastassiadis K, Ang S-L, Stewart AF. 2006. Multiple epigenetic maintenance factors implicated by the loss of Mll2 in mouse development. Dev Camb Engl 133: 1423–1432.

Golemis EA, Brent R. 1992. Fused protein domains inhibit DNA binding by LexA. Mol Cell Biol 12: 3006–3014.

Greer EL, Maures TJ, Hauswirth AG, Green EM, Leeman DS, Maro GS, Han S, Banko MR, Gozani O, Brunet A. 2010. Members of the H3K4 trimethylation complex regulate lifespan in a germline-dependent manner in C. elegans. Nature 466: 383–387.

Halder D, Lee C-H, Hyun JY, Chang G-E, Cheong E, Shin I. 2017. Suppression of Sin3A activity promotes differentiation of pluripotent cells into functional neurons. Sci Rep 7: 44818.

Hallson G, Hollebakken RE, Li T, Syrzycka M, Kim I, Cotsworth S, Fitzpatrick KA, Sinclair DAR, Honda BM. 2012. dSet1 is the main H3K4 di- and tri-methyltransferase throughout Drosophila development. Genetics 190: 91–100.

Han S, Schroeder EA, Silva-García CG, Hebestreit K, Mair WB, Brunet A. 2017. Mono-unsaturated fatty acids link H3K4me3 modifiers to C. elegans lifespan. Nature 544: 185–190.

Hedgecock EM, White JG. 1985. Polyploid tissues in the nematode Caenorhabditis elegans. Dev Biol 107: 128–133.

Heintzman ND, Stuart RK, Hon G, Fu Y, Ching CW, Hawkins RD, Barrera LO, Van Calcar S, Qu C, Ching KA, et al. 2007. Distinct and predictive chromatin signatures of transcriptional promoters and enhancers in the human genome. Nat Genet 39: 311–318.

Ho JWK, Jung YL, Liu T, Alver BH, Lee S, Ikegami K, Sohn K-A, Minoda A, Tolstorukov MY, Appert A, et al. 2014. Comparative analysis of metazoan chromatin organization. Nature 512: 449–452.

Hoe M, Nicholas HR. 2014. Evidence of a MOF histone acetyltransferase-containing NSL complex in C. elegans. Worm 3: e982967.

Howe FS, Fischl H, Murray SC, Mellor J. 2017. Is H3K4me3 instructive for transcription activation? BioEssays News Rev Mol Cell Dev Biol 39: 1–12.

Hu D, Gao X, Morgan MA, Herz H-M, Smith ER, Shilatifard A. 2013. The MLL3/MLL4 branches of the COMPASS family function as major histone H3K4 monomethylases at enhancers. Mol Cell Biol 33: 4745–4754.

Huang C, Yang F, Zhang Z, Zhang J, Cai G, Li L, Zheng Y, Chen S, Xi R, Zhu B. 2017. Mrg15 stimulates Ash1 H3K36 methyltransferase activity and facilitates Ash1 Trithorax group protein function in Drosophila. Nat Commun 8: 1649.

Hughes CM, Rozenblatt-Rosen O, Milne TA, Copeland TD, Levine SS, Lee JC, Hayes DN, Shanmugam KS, Bhattacharjee A, Biondi CA, et al. 2004. Menin associates with a trithorax family histone methyltransferase complex and with the hoxc8 locus. Mol Cell 13: 587–597.

Iwamori N, Tominaga K, Sato T, Riehle K, Iwamori T, Ohkawa Y, Coarfa C, Ono E, Matzuk MM. 2016. MRG15 is required for pre-mRNA splicing and spermatogenesis. Proc Natl Acad Sci U S A 113: E5408–5415.

Jelinic P, Pellegrino J, David G. 2011. A novel mammalian complex containing Sin3B mitigates histone acetylation and RNA polymerase II progression within transcribed loci. Mol Cell Biol 31: 54–62.

Kadamb R, Mittal S, Bansal N, Batra H, Saluja D. 2013. Sin3: Insight into its transcription regulatory functions. Eur J Cell Biol 92: 237–246.

Kasten MM, Dorland S, Stillman DJ. 1997. A large protein complex containing the yeast Sin3p and Rpd3p transcriptional regulators. Mol Cell Biol 17: 4852–4858.

Kelly RDW, Cowley SM. 2013. The physiological roles of histone deacetylase (HDAC) 1 and 2: complex co-stars with multiple leading parts. Biochem Soc Trans 41: 741–749.

Kent WJ, Zweig a S, Barber G, Hinrichs a S, Karolchik D. 2010. BigWig and BigBed: enabling browsing of large distributed datasets. Bioinforma Oxf Engl 26: 2204–7.

Kim T, Buratowski S. 2009. Dimethylation of H3K4 by Set1 recruits the Set3 histone deacetylase complex to 5’ transcribed regions. Cell 137: 259–72.

Laherty CD, Yang WM, Sun JM, Davie JR, Seto E, Eisenman RN. 1997. Histone deacetylases associated with the mSin3 corepressor mediate mad transcriptional repression. Cell 89: 349–356.

Lechner T, Carrozza MJ, Yu Y, Grant PA, Eberharter A, Vannier D, Brosch G, Stillman DJ, Shore D, Workman JL. 2000. Sds3 (suppressor of defective silencing 3) is an integral component of the yeast Sin3[middle dot]Rpd3 histone deacetylase complex and is required for histone deacetylase activity. J Biol Chem 275: 40961–40966.

Lee J, Saha PK, Yang Q-H, Lee S, Park JY, Suh Y, Lee S-K, Chan L, Roeder RG, Lee JW. 2008. Targeted inactivation of MLL3 histone H3-Lys-4 methyltransferase activity in the mouse reveals vital roles for MLL3 in adipogenesis. Proc Natl Acad Sci U S A 105: 19229–19234.

Lee JH, Skalnik DG. 2005. CpG-binding protein (CXXC finger protein 1) is a component of the mammalian Set1 histone H3-Lys4 methyltransferase complex, the analogue of the yeast Set1/COMPASS complex. J Biol Chem 280: 41725–31.

Lee J-H, Skalnik DG. 2008. Wdr82 Is a C-Terminal Domain-Binding Protein That Recruits the Setd1A Histone H3-Lys4 Methyltransferase Complex to Transcription Start Sites of Transcribed Human Genes. Mol Cell Biol 28: 609–618.

Lee J-H, Tate CM, You J-S, Skalnik DG. 2007. Identification and Characterization of the Human Set1B Histone H3-Lys4 Methyltransferase Complex. J Biol Chem 282: 13419–13428.

Lee S, Lee D-K, Dou Y, Lee J, Lee B, Kwak E, Kong Y-Y, Lee S-K, Roeder RG, Lee JW. 2006. Coactivator as a target gene specificity determinant for histone H3 lysine 4 methyltransferases. Proc Natl Acad Sci U S A 103: 15392–15397.

Lenstra TL, Benschop JJ, Kim T, Schulze JM, Brabers NA, Margaritis T, van de Pasch LA, van Heesch SA, Brok MO, Groot Koerkamp MJ, et al. 2011. The specificity and topology of chromatin interaction pathways in yeast. Mol Cell 42: 536–49.

Lewis MJ, Liu J, Libby EF, Lee M, Crawford NPS, Hurst DR. 2016. SIN3A and SIN3B differentially regulate breast cancer metastasis. Oncotarget 7: 78713–78725.

Li H, Durbin R. 2010. Fast and accurate long-read alignment with Burrows-Wheeler transform. Bioinforma Oxf Engl 26: 589–595.

Li T, Kelly WG. 2011. A Role for Set1/MLL-Related Components in Epigenetic Regulation of the Caenorhabditis elegans Germ Line ed. D. Schübeler. PLoS Genet 7: e1001349.

Liu M, Pile LA. 2016. The Transcriptional Corepressor SIN3 Directly Regulates Genes Involved in Methionine Catabolism and Affects Histone Methylation, Linking Epigenetics and Metabolism. J Biol Chem jbc.M116.749754.

Mahadevan J, Skalnik DG. 2016. Efficient differentiation of murine embryonic stem cells requires the binding of CXXC finger protein 1 to DNA or methylated histone H3-Lys4. Gene 594: 1–9.

Margaritis T, Oreal V, Brabers N, Maestroni L, Vitaliano-Prunier A, Benschop JJ, van Hooff S, van Leenen D, Dargemont C, Geli V, et al. 2012. Two distinct repressive mechanisms for histone 3 lysine 4 methylation through promoting 3’-end antisense transcription. PLoS Genet 8: e1002952.

McMurchy AN, Stempor P, Gaarenstroom T, Wysolmerski B, Dong Y, Aussianikava D, Appert A, Huang N, Kolasinska-Zwierz P, Sapetschnig A, et al. 2017. A team of heterochromatin factors collaborates with small RNA pathways to combat repetitive elements and germline stress. eLife 6.

Miller T, Krogan NJ, Dover J, Erdjument-Bromage H, Tempst P, Johnston M, Greenblatt JF, Shilatifard A. 2001. COMPASS: a complex of proteins associated with a trithorax-related SET domain protein. Proc Natl Acad Sci 98: 12902–12907.

Moshkin YM, Kan TW, Goodfellow H, Bezstarosti K, Maeda RK, Pilyugin M, Karch F, Bray SJ, Demmers JAA, Verrijzer CP. 2009. Histone chaperones ASF1 and NAP1 differentially modulate removal of active histone marks by LID-RPD3 complexes during NOTCH silencing. Mol Cell 35: 782–793.

Narayanan A, Ruyechan WT, Kristie TM. 2007. The coactivator host cell factor-1 mediates Set1 and MLL1 H3K4 trimethylation at herpesvirus immediate early promoters for initiation of infection. Proc Natl Acad Sci U S A 104: 10835–10840.

Passannante M, Marti C-O, Pfefferli C, Moroni PS, Kaeser-Pebernard S, Puoti A, Hunziker P, Wicky C, Müller F. 2010. Different Mi-2 complexes for various developmental functions in Caenorhabditis elegans. PloS One 5: e13681.

Patel A, Dharmarajan V, Vought VE, Cosgrove MS. 2009. On the mechanism of multiple lysine methylation by the human mixed lineage leukemia protein-1 (MLL1) core complex. J Biol Chem 284: 24242–56.

Patel A, Vought VE, Dharmarajan V, Cosgrove MS. 2011. A novel non-set domain multi-subunit methyltransferase required for sequential nucleosomal histone H3 methylation by the mixed lineage leukemia protein-1 (MLL1) core complex. J Biol Chem. http://www.ncbi.nlm.nih.gov/entrez/query.fcgi?cmd=Retrieve&db=PubMed&dopt=Citation&list_uids=21106533.

Raja SJ, Charapitsa I, Conrad T, Vaquerizas JM, Gebhardt P, Holz H, Kadlec J, Fraterman S, Luscombe NM, Akhtar A. 2010. The nonspecific lethal complex is a transcriptional regulator in Drosophila. Mol Cell 38: 827–841.

Ramakrishnan S, Pokhrel S, Palani S, Pflueger C, Parnell TJ, Cairns BR, Bhaskara S, Chandrasekharan MB. 2016. Counteracting H3K4 methylation modulators Set1 and Jhd2 co-regulate chromatin dynamics and gene transcription. Nat Commun 7: 11949.

Robert VJ, Mercier MG, Bedet C, Janczarski S, Merlet J, Garvis S, Ciosk R, Palladino F. 2014. The SET-2/SET1 Histone H3K4 Methyltransferase Maintains Pluripotency in the Caenorhabditis elegans Germline. Cell Rep 9: 443–450.

Roguev A, Schaft D, Shevchenko A, Pijnappel WW, Wilm M, Aasland R, Stewart AF. 2001. The Saccharomyces cerevisiae Set1 complex includes an Ash2 homologue and methylates histone 3 lysine 4. EMBO J 20: 7137–7148.

Saha N, Liu M, Gajan A, Pile LA. 2016. Genome-wide studies reveal novel and distinct biological pathways regulated by SIN3 isoforms. BMC Genomics 17. http://www.biomedcentral.com/1471-2164/17/111 (Accessed July 22, 2016).

Sahu SC, Swanson KA, Kang RS, Huang K, Brubaker K, Ratcliff K, Radhakrishnan I. 2008. Conserved Themes in Target Recognition by the PAH1 and PAH2 Domains of the Sin3 Transcriptional Corepressor. J Mol Biol 375: 1444–1456.

Sardiu ME, Smith KT, Groppe BD, Gilmore JM, Saraf A, Egidy R, Peak A, Seidel CW, Florens L, Workman JL, et al. 2014. Suberoylanilide hydroxamic acid (SAHA)-induced dynamics of a human histone deacetylase protein interaction network. Mol Cell Proteomics MCP 13: 3114–3125.

Saunders A, Huang X, Fidalgo M, Reimer MH, Faiola F, Ding J, Sánchez-Priego C, Guallar D, Sáenz C, Li D, et al. 2017. The SIN3A/HDAC Corepressor Complex Functionally Cooperates with NANOG to Promote Pluripotency. Cell Rep 18: 1713–1726.

Schneider R, Bannister AJ, Myers FA, Thorne AW, Crane-Robinson C, Kouzarides T. 2004. Histone H3 lysine 4 methylation patterns in higher eukaryotic genes. Nat Cell Biol 6: 73–77.

Schreiber G, Fersht AR. 1995. Energetics of protein-protein interactions: analysis of the barnase-barstar interface by single mutations and double mutant cycles. J Mol Biol 248: 478–486.

Shilatifard A. 2012. The COMPASS Family of Histone H3K4 Methylases: Mechanisms of Regulation in Development and Disease Pathogenesis. Annu Rev Biochem 81: 65–95.

Shinsky SA, Monteith KE, Viggiano S, Cosgrove MS. 2015. Biochemical Reconstitution and Phylogenetic Comparison of Human SET1 Family Core Complexes Involved in Histone Methylation. J Biol Chem 290: 6361–6375.

Simonet T, Dulermo R, Schott S, Palladino F. 2007. Antagonistic functions of SET-2/SET1 and HPL/HP1 proteins in C. elegans development. Dev Biol 312: 367–383.

Smith HF, Roberts MA, Nguyen HQ, Peterson M, Hartl TA, Wang X-J, Klebba JE, Rogers GC, Bosco G. 2013. Maintenance of interphase chromosome compaction and homolog pairing in Drosophila is regulated by the condensin cap-h2 and its partner Mrg15. Genetics 195: 127–146.

South PF, Fingerman IM, Mersman DP, Du H-N, Briggs SD. 2010. A conserved interaction between the SDI domain of Bre2 and the Dpy-30 domain of Sdc1 is required for histone methylation and gene expression. J Biol Chem 285: 595–607.

Spain MM, Caruso JA, Swaminathan A, Pile LA. 2010a. Drosophila SIN3 Isoforms Interact with Distinct Proteins and Have Unique Biological Functions. J Biol Chem 285: 27457–27467.

Spain MM, Caruso JA, Swaminathan A, Pile LA. 2010b. Drosophila SIN3 isoforms interact with distinct proteins and have unique biological functions. J Biol Chem 285: 27457–27467.

Stempor P, Ahringer J. 2016. SeqPlots - Interactive software for exploratory data analyses, pattern discovery and visualization in genomics. Wellcome Open Res 1: 14.

Steward MM, Lee J-S, O’Donovan A, Wyatt M, Bernstein BE, Shilatifard A. 2006a. Molecular regulation of H3K4 trimethylation by ASH2L, a shared subunit of MLL complexes. Nat Struct Mol Biol 13: 852–854.

Steward MM, Lee JS, O’Donovan A, Wyatt M, Bernstein BE, Shilatifard A. 2006b. Molecular regulation of H3K4 trimethylation by ASH2L, a shared subunit of MLL complexes. Nat Struct Mol Biol 13: 852–4.

Suganuma T, Gutiérrez JL, Li B, Florens L, Swanson SK, Washburn MP, Abmayr SM, Workman JL. 2008. ATAC is a double histone acetyltransferase complex that stimulates nucleosome sliding. Nat Struct Mol Biol 15: 364–372.

Takahashi Y-h., Westfield GH, Oleskie AN, Trievel RC, Shilatifard A, Skiniotis G. 2011. Structural analysis of the core COMPASS family of histone H3K4 methylases from yeast to human. Proc Natl Acad Sci 108: 20526–20531.

Tate CM, Lee J-H, Skalnik DG. 2009. CXXC Finger Protein 1 Contains Redundant Functional Domains That Support Embryonic Stem Cell Cytosine Methylation, Histone Methylation, and Differentiation. Mol Cell Biol 29: 3817–3831.

Thomson JP, Skene PJ, Selfridge J, Clouaire T, Guy J, Webb S, Kerr ARW, Deaton A, Andrews R, James KD, et al. 2010. CpG islands influence chromatin structure via the CpG-binding protein Cfp1. Nature 464: 1082–1086.

Thorvaldsdóttir H, Robinson JT, Mesirov JP. 2013. Integrative Genomics Viewer (IGV): high-performance genomics data visualization and exploration. Brief Bioinform 14: 178–192.

Tyagi S, Chabes AL, Wysocka J, Herr W. 2007. E2F activation of S phase promoters via association with HCF-1 and the MLL family of histone H3K4 methyltransferases. Mol Cell 27: 107–119.

van Nuland R, Smits AH, Pallaki P, Jansen PW, Vermeulen M, Timmers HT. 2013a. Quantitative dissection and stoichiometry determination of the human SET1/MLL histone methyltransferase complexes. Mol Cell Biol 33: 2067–77.

van Nuland R, Smits AH, Pallaki P, Jansen PWTC, Vermeulen M, Timmers HTM. 2013b. Quantitative Dissection and Stoichiometry Determination of the Human SET1/MLL Histone Methyltransferase Complexes. Mol Cell Biol 33: 2067–2077.

van Oevelen C, Bowman C, Pellegrino J, Asp P, Cheng J, Parisi F, Micsinai M, Kluger Y, Chu A, Blais A, et al. 2010. The mammalian Sin3 proteins are required for muscle development and sarcomere specification. Mol Cell Biol 30: 5686–5697.

van Oevelen C, Wang J, Asp P, Yan Q, Kaelin WG, Kluger Y, Dynlacht BD. 2008. A Role for Mammalian Sin3 in Permanent Gene Silencing. Mol Cell 32: 359–370.

Vandamme J, Lettier G, Sidoli S, Schiavi ED, Jensen ON, Salcini AE. 2012. The C. elegans H3K27 Demethylase UTX-1 Is Essential for Normal Development, Independent of Its Enzymatic Activity. PLOS Genet 8: e1002647.

Vermeulen M, Eberl HC, Matarese F, Marks H, Denissov S, Butter F, Lee KK, Olsen JV, Hyman AA, Stunnenberg HG, et al. 2010. Quantitative Interaction Proteomics and Genome-wide Profiling of Epigenetic Histone Marks and Their Readers. Cell 142: 967–980.

Voo KS, Carlone DL, Jacobsen BM, Flodin A, Skalnik DG. 2000. Cloning of a mammalian transcriptional activator that binds unmethylated CpG motifs and shares a CXXC domain with DNA methyltransferase, human trithorax, and methyl-CpG binding domain protein 1. Mol Cell Biol 20: 2108–2121.

Wang P, Lin C, Smith ER, Guo H, Sanderson BW, Wu M, Gogol M, Alexander T, Seidel C, Wiedemann LM, et al. 2009a. Global Analysis of H3K4 Methylation Defines MLL Family Member Targets and Points to a Role for MLL1-Mediated H3K4 Methylation in the Regulation of Transcriptional Initiation by RNA Polymerase II. Mol Cell Biol 29: 6074–6085.

Wang X, Lou Z, Dong X, Yang W, Peng Y, Yin B, Gong Y, Yuan J, Zhou W, Bartlam M, et al. 2009b. Crystal structure of the C-terminal domain of human DPY-30-like protein: A component of the histone methyltransferase complex. J Mol Biol 390: 530–537.

Weiner A, Chen HV, Liu CL, Rahat A, Klien A, Soares L, Gudipati M, Pfeffner J, Regev A, Buratowski S, et al. 2012. Systematic Dissection of Roles for Chromatin Regulators in a Yeast Stress Response. PLOS Biol 10: e1001369.

Wu M, Wang PF, Lee JS, Martin-Brown S, Florens L, Washburn M, Shilatifard A. 2008. Molecular regulation of H3K4 trimethylation by Wdr82, a component of human Set1/COMPASS. Mol Cell Biol 28: 7337–7344.

Wysocka J, Myers MP, Laherty CD, Eisenman RN, Herr W. 2003. Human Sin3 deacetylase and trithorax-related Set1/Ash2 histone H3-K4 methyltransferase are tethered together selectively by the cell-proliferation factor HCF-1. Genes Dev 17: 896–911.

Xiao Y, Bedet C, Robert VJP, Simonet T, Dunkelbarger S, Rakotomalala C, Soete G, Korswagen HC, Strome S, Palladino F. 2011. Caenorhabditis elegans chromatin-associated proteins SET-2 and ASH-2 are differentially required for histone H3 Lys 4 methylation in embryos and adult germ cells. Proc Natl Acad Sci 108: 8305–8310.

Xu L, Strome S. 2001. Depletion of a novel SET-domain protein enhances the sterility of mes-3 and mes-4 mutants of Caenorhabditis elegans. Genetics 159: 1019–1029.

Yao C, Carraro G, Konda B, Guan X, Mizuno T, Chiba N, Kostelny M, Kurkciyan A, David G, McQualter JL, et al. 2017. Sin3a regulates epithelial progenitor cell fate during lung development. Dev Camb Engl 144: 2618–2628.

Yochum GS, Ayer DE. 2002. Role for the mortality factors MORF4, MRGX, and MRG15 in transcriptional repression via associations with Pf1, mSin3A, and Transducin-Like Enhancer of Split. Mol Cell Biol 22: 7868–7876.

Yu BD, Hess JL, Horning SE, Brown GA, Korsmeyer SJ. 1995. Altered Hox expression and segmental identity in Mll-mutant mice. Nature 378: 505–508.

Yücel D, Hoe M, Llamosas E, Kant S, Jamieson C, Young PA, Crossley M, Nicholas HR. 2014. SUMV-1 antagonizes the activity of synthetic multivulva genes in Caenorhabditis elegans. Dev Biol 392: 266–282.

Zargar Z, Tyagi S. 2012. Role of host cell factor-1 in cell cycle regulation. Transcription 3: 187–192.

Zhao X, Su J, Wang F, Liu D, Ding J, Yang Y, Conaway JW, Conaway RC, Cao L, Wu D, et al. 2013a. Crosstalk between NSL Histone Acetyltransferase and MLL/SET Complexes: NSL Complex Functions in Promoting Histone H3K4 Di-Methylation Activity by MLL/SET Complexes ed. P.M. Vertino. PLoS Genet 9: e1003940.

Zhao X, Su J, Wang F, Liu D, Ding J, Yang Y, Conaway JW, Conaway RC, Cao L, Wu D, et al. 2013b. Crosstalk between NSL Histone Acetyltransferase and MLL/SET Complexes: NSL Complex Functions in Promoting Histone H3K4 Di-Methylation Activity by MLL/SET Complexes ed. P.M. Vertino. PLoS Genet 9: e1003940.

